# circDesign Algorithm for Designing Synthetic Circular RNA

**DOI:** 10.1101/2023.07.09.548293

**Authors:** Congcong Xu, Chengtao Pu, Ruofan Chen, Weiyun Wang, Fan Jiang, Changchang Deng, Dongqing Zhai, Yuenan Chen, Weiwei Hu, Yuting Zhang, Yuying Tang, Qiuhe Wang, Jinqi An, He Wang, Jichuan Wu, Xiaotian Wang, Ming Liu, Haifa Shen, Liang Huang, Zhiyuan Zhong, Weihong Tan, Dongsheng Liu, Liang Zhang

## Abstract

Synthetic circular RNA (circRNA) has emerged as a promising platform for vaccine and therapeutic development, featuring its uniqueness in a closed-loop structure, cap-independent translation mechanism, and prolonged expression. However, the rational design of a circRNA sequence to jointly improve its stability and protein coding potential remains challenging. In this study, we present circDesign, an efficient algorithm to achieve the optimal design of circRNA by ensuring optimized folding of each segment, which leads to enhanced circularization efficiency, stability, and translatability. Using rabies virus glycoprotein (RABV-G) as the model antigen, we demonstrated that circDesign-generated circRNAs exhibited higher stability and protein translation efficiency *in vitro* and *in vivo* compared to other codon adaptation index (CAI)-optimized sequences, thus leading to enhanced *in vivo* immunogenicity. Ribosome and polysome profiling further revealed that an intact internal ribosome entry site (IRES) structure is critical for efficient translation. By intentionally disrupting the IRES motifs, we observed that the resulting circRNA sequences had lower translatability compared to circDesign-generated sequences. Taken together, our circular RNA design algorithm provides a general strategy to leverage the capability of circRNA as next-generation vaccines or therapeutics.

## Introduction

Synthetic circular RNA (circRNA) is emerging as next-generation RNA platform for vaccine or therapeutic development^1,2,3^. Distinct from conventional linear mRNA, circRNA is featured with a closed-loop structure, rendering it a unique biomolecule in terms of synthesis, stability, and translation initiation mechanism ^4,5^. Therefore, circRNA sequence engineering strategy differs from a simplified linear mRNA design approach from the following aspects. Firstly, *in vitro* circRNA synthesis process involves circularization step which typically adopts the catalytic activity from ribozyme sequence for sequence ligation, as exemplified by group I/II intron-mediated self-splicing method or *trans*-ribozyme-based circularization (TRIC) ^6,7,8,9^. Secondly, the resultant synthetic circRNA with an endless structure exhibited a different stability profile compared with linear mRNA owing to its exonuclease resistance and different biological decay mechanisms ^10^. Lastly, synthetic circRNA typically adopts viral internal ribosome entry site (IRES) motif for cap-independent translation initiation, which is generally not as efficient as *N* ^7^-methylguanosine (m^7^G) cap present on linear mRNA for the translation initiation ^11^. Taken together, achieving optimal circular RNA sequence design necessitates delicate consideration of circRNA folding which governs circularization, stability, and translation initiation.

Recent studies highlighted the importance of augmenting translation initiation of circRNA for improved protein coding capacity. Different approaches such as introducing branched or internal cap modification, eIF4G-recruiting aptamer, or IRES-like element have been proposed to overcome the inherent limitations of IRES scaffold-mediated initiation efficiency ^12,13,14^. Despite of these advancements, there remains an urgent demand for a rational circRNA engineering paradigm that synergistically achieves three critical objectives: 1) preservation of highly efficient circularization; 2) structural compactness of circRNA for enhanced stability; 3) full exploitation of IRES-mediated initiation capacity by maintaining its structural integrity. The development of such an integrated design framework represents a crucial unmet need in the field of synthetic circRNA therapeutics.

We have previously reported LinearDesign algorithm for designing coding sequence (CDS) region of linear mRNA ^15^. This modular optimization approach by dissecting untranslated region (UTR) and CDS is practically feasible for conventional linear mRNA due to the simplified molecular structure where short UTR region (∼ 100 nt in length) has negligible structural interference with the stringent folding of long CDS region. Therefore, a brutal combination of independently generated CDS with UTR sets led to improved translation efficiency, and *vice versa* ^16,17^. However, the sequence design for circRNA is further complicated by the long IRES motifs which necessitate correct folding for efficient recruitment of translational machineries^18,19,20^.

Meanwhile, the secondary structure dictated by primary sequence affects circularization efficiency and stability as well. Therefore, a rational circRNA sequence design algorithm aims to address these issues by taking account of circRNA molecule as an entirety instead of segmented modules.

Here we present an efficient algorithm called **circDesign** for designing circRNA with improved circularization efficiency, stability, and translatability. In this approach, given a specific IRES sequence for translation efficiency and catalytic region for circularization, the CDS region is jointly generated within the entire design space, integrating minimum free energy (MFE), codon adaptation index (CAI), and IRES structural deviation. MFE value dictates the compactness of circRNA structure and CAI of CDS refers to the codon optimality as previously reported ^15^. Specifically, we introduced a constraint parameter, IRES structural deviation, which quantifies the structural specificity required by the IRES region to minimize interference with other regions, particularly the CDS (Fig. 1a). Using rabies virus glycoprotein (RABV-G) as a model antigen, we utilized circDesign algorithm to generate optimal circRNA sequences. Compared with other conventionally generated circRNA, sequences generated with circDesign demonstrated improved circularization efficiency, stability, protein translatability as well as *in vivo* immunogenicity. Further ribosome and polysome profiling results revealed the importance of IRES structural integrity in circRNA translation as well as biological stability. We believe the circDesign approach provides a general strategy for synthetic circular RNA design and holds immense potentials for broader circRNA-based vaccine and therapeutic development.

**Figure 1.**
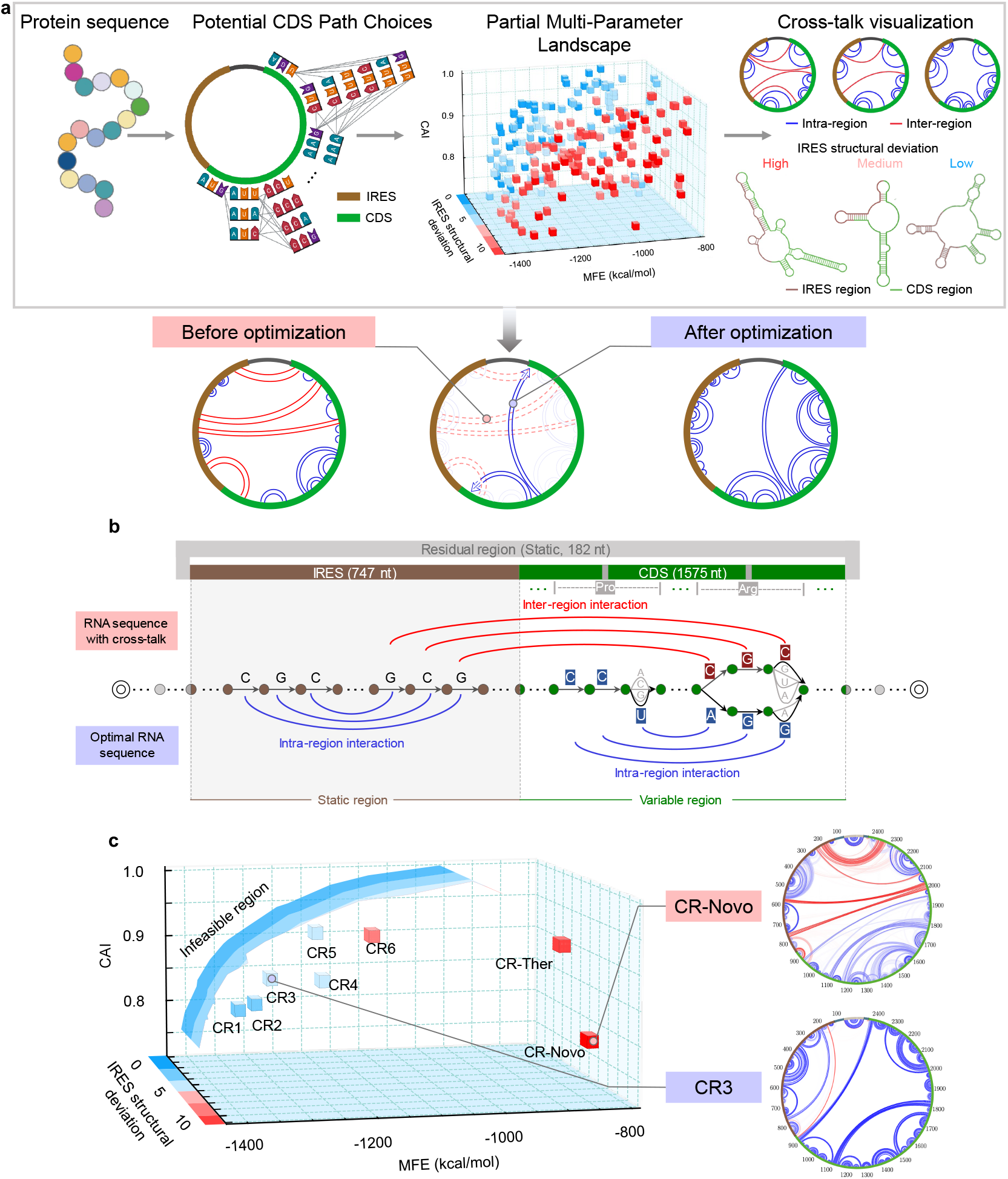
circDesign workflow using RABV-G as a model antigen. **a**, Schematic representation of the circDesign workflow for generating optimized circRNA constructs. For a given protein sequence, circDesign explores potential CDS path choices with three key parameters: Minimum Free Energy (MFE), Codon Adaptation Index (CAI), and IRES structural deviation. Cross-talk visualization: In the circular plot, red lines represent undesirable interactions between the IRES and other regions, while blue lines denote base pairing within the same region. **b**, Detailed illustration of the circDesign algorithm’s optimization process. Codon choices that lead to undesirable interactions (depicted as red lines) are penalized, while those that enhance stable internal structures within each region (depicted as blue lines) are favored. **c**, Visualization of circDesign-generated sequences alongside two benchmark sequences within the search space defined by MFE, CAI, and IRES structural deviation. Base-pairing probabilities are illustrated for the optimized sequence CR3 and the benchmark sequence CR-Novo. In the circular plot, the darkness of each line reflects the strength of the pairing probability

## Results

### circDesign algorithm for circular RNA design

We developed circDesign to enable rational circRNA design for next-generation therapeutics. Unlike linear mRNA, which relies on short UTRs for translation initiation, circRNAs typically include long viral or cellular IRES sequences (500–800 nucleotides) that must independently fold into a hierarchical structure to efficiently recruit the translational machinery. A key challenge is minimizing “cross-talk” between the CDS (typically the longest region in the overall circRNA sequence) and IRES regions ^20,21,22^. To address this, circDesign employs dynamic programming to jointly optimize mRNA stability (measured by MFE), codon optimality (measured by CAI), and the structural integrity of the IRES (measured by IRES structural deviation). During the dynamic programming process, an explicit penalty is applied to reduce unwanted interactions between the IRES and CDS (Fig.1b, red lines) while promoting favorable internal secondary structures (Fig.1b, blue lines).

Furthermore, circDesign introduces a post-processing step to address the variability in residual sequences arising from different circularization methods and the incorporation of additional elements for specific objectives. The full circRNA sequence, including these variable regions, is assembled, and the IRES structural deviation is calculated by comparing the predicted IRES structure within the full circRNA to that of the standalone form. Under similar MFE and CAI conditions, the sequence with the minimal IRES deviation is selected. (See Methods, “Overall algorithmic rationale and innovation” for further details.)

### Design, synthesis, and characterization of circular RNA encoding RABV-G

Herewith RABV-G encoding circRNA, we adopted IRES from Coxsackievirus B3 (CVB3) and applied circDesign to generate six distinct sequences (CR1–6) alongside two benchmark sequences for comparison ^23^. The benchmark sequences, CR-Ther and CR-Novo, were designed using publicly available platforms from Thermo Fisher and Novoprotein, respectively. These platforms primarily optimize codon preference, but neither account for secondary structure nor ensure the integrity of the structure of IRES. In contrast, the six circDesign-generated sequences are strategically distributed within low MFE regions (MFE *< −* 1100 kcal*/*mol). Among them, CR1 and CR2 exhibit similar CAI with CR-Novo, while CR5 and CR6 have CAI values comparable to CR-Ther. Additionally, CR3 and CR4 were designed as MFE-balanced complements to CR2 and CR5, respectively, maintaining similar MFE levels while achieving higher CAI values. Finally, compared to the benchmark sequences, CR1–CR6 demonstrate a 1.5-to 4.5-fold improvement in IRES structural deviation, which might induce a better preservation of IRES functionality.

To synthesize the circRNA, circularization was facilitated by the self-cleaving permuted group I catalytic intron as previously reported, enabling two exon fragments (E1 and E2) to join through a pair of constitutive transesterification reactions ^24^. The secondary structures of CR1–6 along with two benchmark sequences were predicted (see Extended Data Fig. 1,2). Taking CR3 and CR-Novo as representative sequences, CR3 was designed with a significantly lower MFE while maintaining a comparable CAI. Additionally, CR3 shows considerably reduced inter-region interaction (depicted as red lines) between CDS and IRES compared to CR-Novo (Fig. 1c). As a result, CR3 forms a more stable secondary structure in both the IRES and CDS regions than CR-Novo. This outcome could be attributed to our circDesign, which intentionally considers and minimizes structural interactions between the IRES and CDS regions during sequence generation. Similar trends were observed in other sequences generated by circDesign (see Extended Data Fig. 1,2).

Different designs exhibited variations not only in MFE, CAI value, and IRES structural deviation, but also demonstrated differences in circularization efficiency. Considering a more compact IRES and CDS folding which might lead to less interference with the formation of active intron, the circDesign-generated sequences (CR1–6) displayed relatively higher circularization efficiencies compared to CR-Novo and CR-Ther (Fig. 2a). To remove impurities from the circularization reaction products, size-exclusion column chromatography was used to obtain purified circRNA (see Extended Data Fig. 3a) which induced minimized innate immunogenicity (see Extended Data Fig. 3b–c). To characterize the circRNA, non-denaturing agarose gel electrophoresis was performed at room temperature. As shown in Fig. 2b, it is intriguing to observe that the gel mobility of circRNAs was well correlated with their MFE values given their identical nucleotide lengths. The CR-Novo sequence, which exhibited the highest MFE, migrated the slowest in the gel matrix—likely due to its less compact structure—while circDesign-generated sequences, particularly CR1–5, migrated faster as a result of their highly compact folding.. These findings underscore that a fully integrated circRNA design aimed at achieving a more compact overall structure may be key to effectively regulating its folding paradigm.

**Figure 2.**
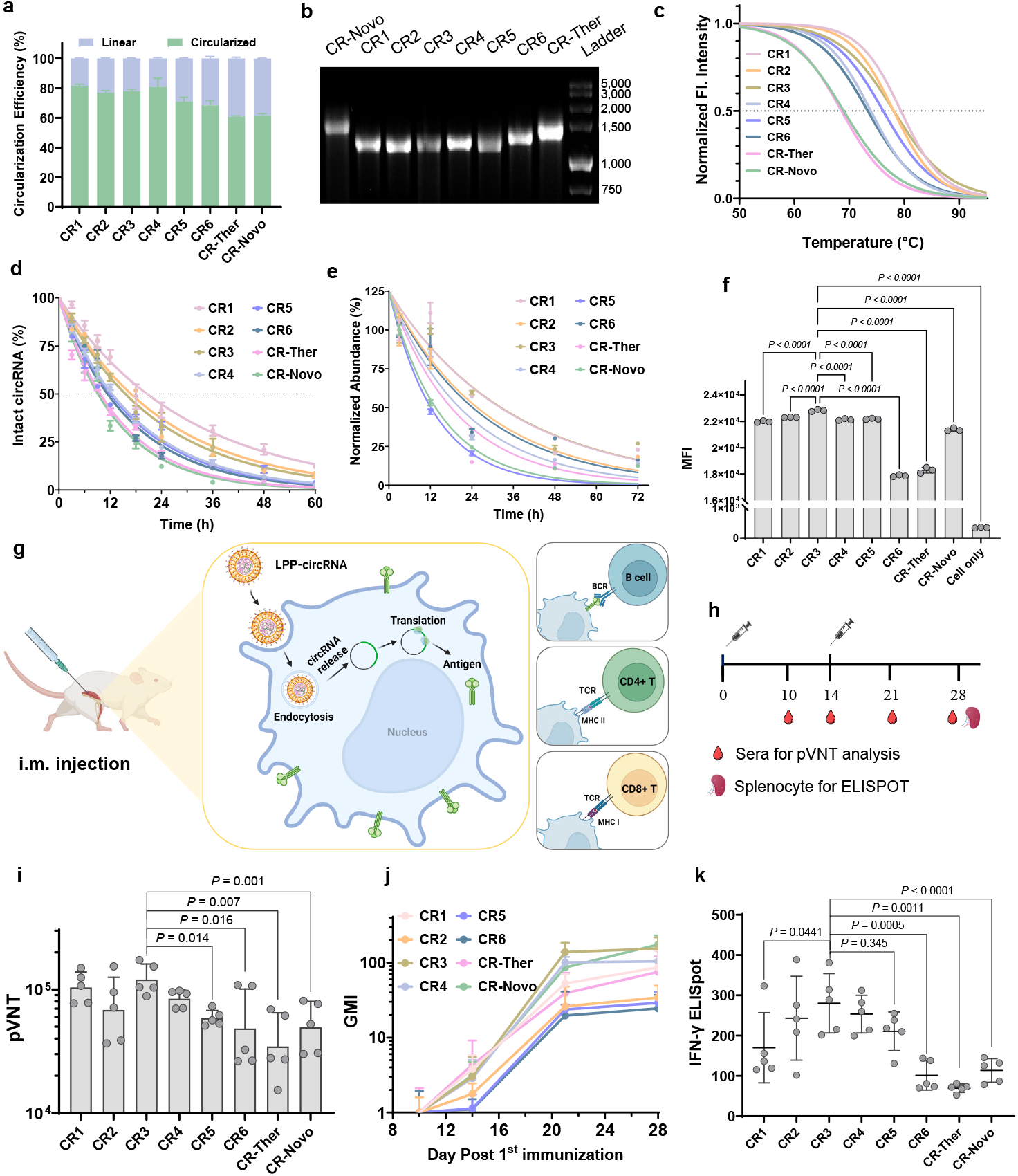
Characterization of circRNAs Encoding RABV-G. **a**, Comparison of the circularization efficiency for circRNAs (CR1–6, CR-Novo, CR-Ther) measured by qPCR. Data are presented as mean ± s.d. for *n* = 3 independent replicates. **b**, Native agarose gel electrophoresis of circRNAs. Faster migration of CR1–5 compared to CR-Novo and CR-Ther reflects greater folding compactness. **c**, Thermodynamic stability analysis of circRNAs, suggesting a correlation of MFE with melting temperature (*T*_m_). *T*_m_ values were extrapolated from sigmoidal curves fitted to the mean values of three independent measurements. **d**, *In vitro* degradation kinetics of circRNAs in PBS buffer containing 10 mM Mg^2+^ at 37 ^*°*^C. Data are presented as mean ± s.d. (*n* = 3) and fitted with a one-phase decay curve to determine half-life and decay constant. **e**, Intracellular degradation kinetics of circRNAs in Human Skeletal Muscle Cells (HSkMC) cells over 72 h post-transfection. Data are presented as mean ± s.d. (*n* = 3) and fitted with a non-linear one-phase decay curve. **f**, Flow cytometry analysis of circRNAs expression in HSkMC 24 h post-transfection. Mean fluorescence intensity (MFI) represents corresponding protein expression levels. Data are presented as mean ± s.d. (*n* = 3). Statistical analysis was performed using one-way ANOVA with multiple comparisons against the CR-Novo group. **g**, Immunization schedule for Balb/c mice (*n* = 5 per group) and proposed mechanism of LPP-formulated circRNA rabies vaccine. **h**, Specific time points for the detection of pseudovirus neutralizing titers (pVNT) in mouse sera and ELISpot analysis of mouse splenocytes. **i**, pVNT in sera from mice immunized with two doses of LPP-formulated RNA vaccines on day 21 post-immunization. **j**, Geometric mean increase (GMI) of pVNT from mice vaccinated with different RNA vaccines, using pVNT on day 10 as the baseline. GMI data are plotted with geometric standard deviation. **k**, IFN-*γ* ELISpot analysis of splenocytes from mice on day 28, demonstrating cellular immune responses triggered by circDesign-generated sequences. Data are represented as spot-forming cells (s.f.c.) per 3 × 10^5^ cells. Each point on the plot represents a single mouse, with the bold black line indicating the median. Statistical significance between groups was determined using one-way ANOVA with Dunn’s multiple comparisons test.

### Generated circRNA exhibited enhanced *in vitro* stability and protein expression

As the secondary structure dictates the functionally half-life of mRNA, we then extensively evaluated the *in vitro* stability of circRNAs in terms of thermodynamic folding, in-solution degradation as well as intracellular decay ^25^. By heating the circRNA in solution to gradually unfold its structure, the fluorescence intensity from RNA bound RiboGreen^®^ reagent attenuated accordingly. We calculated the melting temperature (*T*_m_) of circRNA from the fluorescent intensity curve as Fig. 2c. CR1–3 had slightly higher *T*_m_ than CR4–6, with an obviously increased thermodynamic stability compared to CR-Novo and CR-Ther. The melting curve results was associated with MFE values.

Like linear mRNA, the chemical instability of circRNA originated mainly from hydrolysis reaction despite of its endless structure. To examine the in-solution stability of all circRNAs, RNA samples were incubated in PBS buffer containing 10 mM Mg^2+^ at 37 ^*°*^C mimicking the physiological buffering system over a time course of 60 h. At different time points, the integrity of circRNA was assessed. The degradation of circRNA as a function of time was fitted to a one-phase decay curve, suggesting different degradation kinetics. circRNAs with higher MFE values (CR-Novo & CR-Ther) underwent faster degradation kinetics with half-life (*t*_1*/*2_) of 9.14 h and 9.68 h respectively, while CR1–3 had considerably extended *t*_1*/*2_ ranging from 15.56 h to 20.62 h (Fig. 2d).

Apart from chemical degradation, the intracellular decay of exogenous circRNA is also regulated by different pathways in related to its translational activity ^10,18^. To assess the in-cell half-life of circRNA, the abundance of circRNA extracted from cells was quantified using reverse transcription quantitative real-time PCR (RT-qPCR) method at various time points (3 h, 12 h, 24 h, 48 h and 72 h) post-transfection into Human Skeletal Muscle Cells (HSkMC). As illustrated in Fig. 2e, a slightly different decay pattern of all tested circRNA was observed. CR1 and CR3 had a prolonged intracellular half-life (24.34 h and 19.31 h, respectively) while CR-Novo and CR5 showed a faster in-cell decay (10.21 h and 9.18 h, respectively). As the biological fate of transfected circRNA is associated with a plethora of factors including translation status and nuclease resistance, the in-cell degradation of artificially synthesized circRNA could possibly be controlled by tuning its secondary structure and translation efficiency. We reasoned that CR3 sequence had decent stability in conjunction with high translation efficiency, enabling its resistance against chemical degradation and biological decay.

Since the stability and translation efficiency of circRNA co-regulates the protein output level, we evaluated the RABV-G expression from transfected circRNA in HSkMC using flow cytometry (Fig. 2f). At 24 h post-transfection, antigen was highly expressed in CR3 group as suggested by the mean fluorescent intensity (MFI). CR1–5 had a remarkably higher protein output level than CR6 and CR-Ther, largely attributed to the relatively instable structure of CR6 and CR-Ther. However, it’s interesting to find that benchmark CR-Novo exhibited a decent protein coding capacity despite of its lower CAI value compared to CR-Ther. This evidence emphasized the importance of taking multiple influential factors into consideration other than CAI to stratify the mRNA coding potentials.

### LPP delivery system efficiently delivered circRNA *in vitro* and *in vivo*

To investigate the performance of circRNA *in vivo*, a proprietary lipopolyplex (LPP) delivery system was used to formulate the circular RNA ^26^. As shown in Extended Data Fig. 4a, following a two-step microfluidic mixing process, cationic polymer complexed circRNA could be encapsulated in a bilayer liposome-like nanoparticle with a “core-shell” structure. Previous studies have demonstrated the efficiency and safety of using this LPP delivery system to deliver linear mRNA ^27^. Herewith circRNA encoding GFP (CR-GFP) or firefly luciferase (CR-FLuc) was formulated into LPP to study its *in vitro* cell expression and *in vivo* biodistribution compared to its chemically modified linear counterpart (LR-GFP or LR-FLuc), respectively. Flow cytometry analysis of cells treated with LPP formulated CR-GFP showed a dosage-dependent translation.

Stronger expression was observed in circular RNA group than that of LR-GFP (Extended Data Fig. 4b). Besides, the protection of encapsulated circRNA against RNase digestion was evaluated in LPP and lipid nanoparticle (LNP), a widely used mRNA delivery system. The residual circRNA was quantified at 30 min post-treatment and the results suggested a higher protection efficacy against nuclease digestion using LPP formulation (Extended Data Fig. 4c).

Upon intramuscular administration of LPP formulated CR-FLuc and LR-FLuc in mice, *in vivo* imaging system (IVIS) captured the whole-body *in vivo* luminescence activities at different time points (6 h, 24 h, 48 h, 72 h, 96 h and 168 h) (Extended Data Fig. 4d), reflecting the expression kinetics of luciferase from CR-FLuc or LR-FLuc. It turned out that circular RNA produced luciferase in a more durable manner with higher accumulated expression as shown in Extended Data Fig. 4e. Though with similar physicochemical properties of LPP formulation and lower onset expression as seen at 6 h post administration, CR-FLuc showed extended expression over the course of one week while luminescence from LR-FLuc group descended to background level within a relatively shorter time (2 days post-dosing). A gross estimation of accumulated expression was conducted simply by calculating the area-under-curve (AUC) of total flux of luminescence. CR-FLuc group led to approximately 5.3-fold of total luciferase expression compared to its linear counterpart over the course of 8 days (Extended Data. 4f). Indeed, other group has reported similar translation kinetics of circular RNA compared to linear mRNA, thus rendering circRNA a promising therapeutic or vaccine platform for sustained protein supplement or enduring protection against pathogens. These results suggested the feasibility of using LPP for efficient *in vivo* delivery of circRNA despite of its closed loop feature.

### circDesign-generated circRNA rabies vaccine demonstrated superior *in vitro* stability and *in vivo* performance

It has been reported that linear mRNA encapsulated in lipid nanoparticles still underwent slow transesterification hydrolysis ^28^. In this study, the stability of LPP-circRNA nanoparticles was evaluated to simulate the vaccine storage condition under the formulated situation. Henceforth, the same lipid composition was adopted to encapsulate circDesign-generated CR1–6 and benchmark circRNA encoding RABV-G to rule out the impact from delivery system on their *in vitro* and *in vivo* performance. The *in vitro* characterization of formulated circRNAs showed that LPP nanoparticles had similar size of around 110 nm (Extended Data Fig. 4g) and positive zeta potentials (in the range of 20–35 mV) (Extended Data Fig. 4h) regardless of the encapsulated circRNA. However, the importance of optimizing the stability of circRNAs was manifested in their storage stability in lipid formulation. We examined the mRNA content of LPP-circRNA complex on day 30 at room temperature to assess the integrity of RNA component. A faster RNA degradation in LPP formulated CR-Novo and CR-Ther was observed (Extended Data Fig. 4i) with 15–20% circRNA loss on day 30 while circDesign-generated circRNAs (CR1–5) showed little degradation over tested period.

We then evaluated the *in vivo* immunogenicity of RABV-G encoded RNA vaccines. By intramuscular injection into mice, it’s proposed that the circRNA-LPP nanoparticles entered antigen-presenting cells *via* endocytic pathway and circRNA got released from endosome into cytoplasm for antigen translation. The translated antigens were then processed to elicit humoral and cellular immune responses as illustrated in Fig. 2g. Two doses of rabies vaccines were *i*.*m*. administered at an interval of two weeks in Balb/c mice (*n* = 5 per group). Mice were weighted after first dosing and no weight loss was observed. Mice sera were collected on day 10, 14, 21 & 28 to assess the pseudovirus neutralizing antibody titer (pVNT) and spleens were extracted for cellular immunity assessment on day 28 (Fig. 2h). As neutralizing antibodies (nAbs) play a critical role in protection against rabies virus, monitoring the geometric titer (GMT) nAb level gave us a hint on the efficacy of rabies vaccines ^29^. On day 10 post-priming dose, overall antibody titers remained low across all groups, consistent with an early-stage immune response. However, a distinct pattern emerged among the circular RNA (circRNA) constructs. Specifically, CR1–2 and CR5–6 exhibited higher antibody titers compared to the two benchmark sequences, CR-Novo and CR-Ther, while CR3 and CR4 achieved titers comparable to these benchmarks (Extended Data Fig. 5a). This early divergence hinted at the potential of certain circDesign-generated sequences to induce a more robust humoral response, even at this preliminary stage. By day 14 post-priming, the antibody titer expression closely paralleled the trends observed on day 10 (Extended Data Fig. 5b). One week after booster (day 21), the circDesign-generated sequences (CR1–4) demonstrated a clear advantage in eliciting stronger humoral immune responses compared to the bench-mark circular sequences, CR-Novo and CR-Ther (Fig. 2i). Among the generated sequences, CR3 emerged as a standout performer, exhibiting a 3.5-fold increase in antibody titer relative to CR-Ther and a 2.4-fold increase compared to CR-Novo. All circular RNA groups exhibited continued increase in pVNTs on day 28, CR3 consistently maintained higher antibody titers relative to the two benchmark sequences, cementing its superior performance across the study duration (Extended Data Fig. 5c). The geometric increase (GMI) of pVNT was calculated based on the GMT on day 10 as baseline to show the trend of humoral immune response induction. As shown in Fig. 2j, following a booster vaccination (from day 14 to day 21), all circDesign-generated sequences exhibited faster immune response kinetics compared to CR-Ther. Notably, CR3 and CR4 outperformed CR-Novo and maintained sustained immune responses from day 21 to day 28. The detailed pVNT kinetics for each group over time, presented in (Extended Data Fig. 5d), highlight the significant potential of circDesign enabled vaccine design in eliciting robust antibody responses. It’s also worth noting that cellular immune responses induced by circDesign-generated sequences (CR1–4) especially CR3 are more prominent than other groups (Fig. 2k). A higher frequency of RABV-G specific T cells secreting interferon *γ* (IFN-*γ*) was observed in CR3 group with a mean value of 280.1 spots counted using Enzyme-linked immunosorbent spot (ELISpot) assay (Extended Data Fig. 6). Though the practical significance of higher RABV-G specific cellular immunity in rabies prevention remains elusive, we speculate the improved T cell immune responses from circDesign-generated vaccine may play important roles in rabies virus clearance from the host ^30^. Taking a closer look at the cellular immune activation profile, flow cytometry analysis of the RABV-G peptide stimulated T cells was performed. Following a stepwise gating of the cell population (Extended Data Fig. 7), the ratio of CD4^+^ and CD8^+^ T cells producing IFN-*γ*, TNF-*α*, IL-2 and IL-4 were computed, respectively. The results indicated the strong RABV-G specific CD4^+^ and CD8^+^ T cell responses elicited by circDesign-generated vaccines (Extended Data Fig. 8). We observed higher ratios of IFN-*γ*, TNF-*α* and IL-2 secreting T cells but minimal IL-4 producing population in both CD4^+^ and CD8^+^ T cells (Extended Data Fig. 8), suggesting a *T*_*h*_1-biased immune responses induced by circRNA rabies vaccines.

Combined with the in-solution and in-cell stability results of naked or LPP formulated circRNA, these results convincingly demonstrated the improved potency from circDesign-generated sequences owing to their well-defined structure and enhanced stability during storage and intracellular trafficking.

### Ribosome and polysome profiling revealed the enhanced translation capacity of circDesign-generated circRNA

To elucidate the mechanism underlying the superior *in vitro* and *in vivo* performance of circDesign-generated circRNA, we conducted a side-by-side comparison of CR3 and CR-Novo using ribosome profiling and polysome profiling analyses. Beyond differences in MFE and CAI, secondary structure predictions revealed another key distinction: CR3 maintains an intact structure of IRES, while in CR-Novo, the IRES interacts with the CDS disrupting its structure (Fig. 3a,b). This structural integrity in CR3 likely enhances its translational performance.

**Figure 3.**
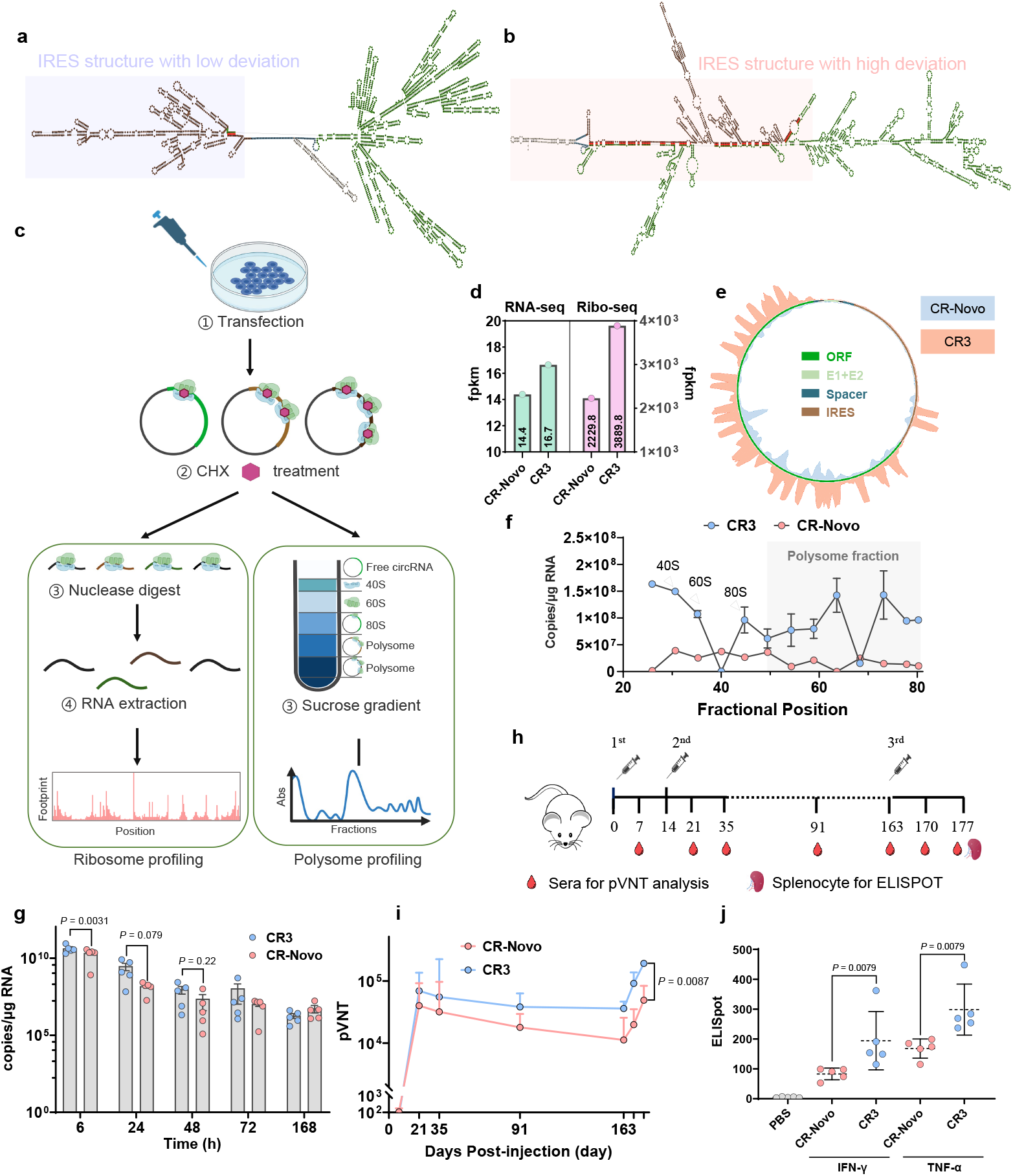
Comparison of Translation Efficiency between CR3 and CR-Novo. **a–b**, Predicted secondary structures of CR3 (a) and CR-Novo (b). **c**, Schematic workflow of ribosome profiling and polysome profiling for circRNA translation analysis. **d**, Bar graph of RNA-seq and Ribo-seq fpkm values for CR3 and CR-Novo, showing higher abundance and translation efficiency in CR3. **e**, Circular bar plot illustrating ribosome occupancy on CR3 and CR-Novo transcripts. **f**, RT-qPCR quantification of CR3 and CR-Novo in sucrose gradient fractions following ultracentrifugation. Arrows indicate the positions of 40S, 60S, 80S (monosome), and polysome fractions are highlighted by shadow. Data are presented as mean ± s.d. (*n* = 3 independent biological replicates). **g**, Time-course comparison of circRNA abundance in muscle tissue. Statistical significance was determined by multiple Mann-Whitney U tests. **h**, Schematic diagram of the immunization schedule in the Balb/c mouse study (*n* = 5). **i**, Kinetics of pseudovirus neutralizing titers (pVNT) for CR3 and CR-Novo immunization. Data are presented as geometric mean ± geometric s.d.. Statistical significance was determined by multiple Mann-Whitney U tests. **j**, IFN-*γ* and TNF-*α* ELISpot assay of splenocytes from immunized mice on Day 28, showing enhanced cellular immune responses elicited by CR3. Data are presented as spot-forming cells (s.f.c.) per 3 × 10^5^ cells. Each data point represents an individual mouse, with the median indicated by a bold black line. Statistical significance was determined using one-way ANOVA with Dunnett’s multiple comparisons test.

After transfecting cells and following the standard workflow (Fig. 3c), ribosome footprints were aligned to full-length CR3 and CR-Novo. The ultracentrifugation fractions were quantified to determine the fractional composition of circRNA. In both RNA direct sequencing counts and ribosome sequencing counts (Fig. 3d), CR3 had higher fpkm (fragment per kilobase of transcript per million mapped reads) of RNA-seq and ribo-seq than that of CR-Novo at the time of cell being harvested, indicating a higher circRNA abundance and ribosome footprints of CR3. The translation efficiency (TE)—calculated as footprint abundance relative to mRNA abundance—was 1.5-fold higher in CR3 (233.7) compared to CR-Novo (159.9), highlighting CR3’s superior protein production capacity. After alignment of the ribosome footprints on circRNA, it’s clear that the cap-independent translation mechanism of circRNA driven by CVB3 IRES efficiently recruits ribosome onto circular RNA to initiate the protein translation. Most of the ribosome protected fragments were identified in CDS region in both CR3 and CR-Novo, though the read count and distribution varied between these two groups. As suggested in Fig. 3e, the ribosomal footprints were mapped and distributed more evenly throughout the CDS region while CR-Novo had more distinct footprint accumulation peaks. The fully aligned ribosomal footprints were plotted on circular CR3 and CR-Novo annotated with each functional region, respectively (Extended Data Fig. 9b,d). In addition, several major peaks upstream of CDS indicated the recruitment of ribosome for translation initiation. Taking a closer look at the secondary structure and pairwise visualization (Extended Data Fig. 9a,c), it’s highly possible that the disrupted IRES motif in CR-Novo impeded the translational machinery recruitment, thus resulting in a much lower ribosome footprint abundance. In addition, polysome profiling was performed on CR3 and CR-Novo to compare the ribosome loading capacity. After ultracentrifugation, total RNA in each fraction was quantified, with the 80S peak representing monosomes and later peaks indicating polysome-bound circRNA. CR3 displayed higher ribosome loading than CR-Novo (Fig. 3f), reinforcing its enhanced translation efficiency.

In summary, CR3 generated by circDesign, preserves the integrity of its IRES structure, enabling efficient ribosome recruitment and translation, which explains its superior performance over CR-Novo.

### The *in vivo* stability of CR3 contributed to durable humoral immune responses

The abovementioned *in vitro* and *in vivo* evaluation of circRNAs implicated the importance of circRNA stability as well as translation efficiency in achieving optimal performance. Therefore, balancing the optimization of CAI (translation efficiency) and MFE (stability), while preserving the integrity of the IRES structure, is essential for creating a well-defined circRNA folding structure. This balanced approach likely contributes synergistically to superior performance, maximizing *in vivo* outcomes. Of all circDesign-generated sequences, CR3 showed superior stability, protein translation output, along with *in vivo* immunogenicity. Compared to CR-Novo, it’s intriguing to find that another benchmark sequence CR-Ther had lower protein translation and induced relatively weaker immune responses despite its higher CAI. Consequently, we further compared CR-Novo with CR3 regarding their *in vivo* stability to explain the advantages of circDesign-generated CR3 over CR-Novo in their potency and durability. To study the *in vivo* stability, LPP formulated CR3 and CR-Novo were i.m. administered into Balb/c mice. At 6 h, 24 h, 48 h, 72 h and 168 h post-dosing, mice were sacrificed and total RNAs were extracted from different organs including muscle (injection site), lymph node, spleen, liver, lung, heart, kidney, brain and small intestine (sm. int.). A standard plasmid containing the target sequence was used as internal standard control to quantify the circRNA copies per µg of total RNA using qPCR analysis. the good safety profile of LPP delivery system, as major injected dose accumulated at the injection site without significant retention in liver both for CR-Novo (Extended Data Fig. 10a) and CR3 (Extended Data Fig. 10b). Moreover, as time went by, more fractions of circRNA were detected in spleen and lymph node where the immune activation mainly occurs. It’s noted that CR3 had extended muscle retention proportion (88.5 %, 70 % and 29.4 % at 24 h, 48 h and 72 h, respectively) compared to CR-Novo (74.5 %, 25.4 % and 7.4 % at 24 h, 48 h and 72 h, respectively) (Extended Data Fig. 10c). It’s hypothesized that higher *in vivo* stability of CR3 benefited the longer retention at injection site given the same dosage as CR-Novo. Within 72 h of administration, the circRNA quantification data of muscle at injection site suggested the higher abundance of CR3 (Fig. 3g). This potentially contributed to a sustained lymphatic migration of antigen presenting cells (APCs) to peripheral immune organs such as draining lymph nodes and spleen, leading to robust immune activation.

Above findings rendered CR3-LPP a more potent rabies vaccine candidate potentially inducing long-lasting adaptive immunity against rabies virus. To study the longevity of immune responses, Balb/c mice were immunized with LPP formulated CR3 and CR-Novo with the same dosage (2 µg per immunization for prime and boost at an interval of two weeks). The mice sera were collected over a much longer period during which the nAb titer was monitored (Fig. 3h). As shown in Fig. 3i, CR3 induced a stronger humoral immune response following prime and booster. 14 days after booster dose (on day 35), the pVNT of CR3 group remained 1.7-fold of CR-Novo (Fig. 3i). With a relatively slower waning rate of pVNT, CR3 group exhibited 2.1-fold and 3.2-fold pVNT over CR-Novo group on day 90 and 163, respectively. According to the guidelines of rabies vaccine released by World Health Organization (WHO), a second booster was recommended when re-exposure to rabies virus occurred 3 months after the last booster. Thus, we administered the second booster in mice on day 163 to evaluate the immune responses after re-boost. The second booster induced much stronger immune response in CR3 group with 4.6-fold and 3.9-fold pVNT of CR-Novo one week (day 170) and two weeks (day 177) after re-booster dose (Fig. 3i). Besides, the cellular immune responses were assessed by ELISpot on day 170 using mice spleenocytes. CR3 immunized mice had shown significantly stronger antigen-specific IFN-*γ* and TNF-*α* secretion (Fig. 3j), potentially leading to better clinic outcome.

### IRES structural integrity impacts circRNA translation and stability

In our circDesign approach, we introduced IRES structural deviation—a measure of IRES structural integrity— as a key parameter to optimize alongside MFE and CAI. In the previous section, we compared CR3 with CR-Novo and observed that CR3, which exhibits greater IRES structural integrity, outperformed CR-Novo in circular RNA performance. However, as CR3 and CR-Novo also differed in MFE and CAI, we could not exclude the influence of these factors on the observed performance. To specifically investigate the intricate relationship between IRES structural integrity and the translational performance of circRNAs, we designed a series of circRNA constructs (CR16111, CR1120, CR3661, CR929, and CR6223) with comparable MFE and CAI but varying degrees of IRES structural deviation using CR3 as a reference sequence (Fig. 4a). Protein expression revealed that CR3, characterized by the lowest IRES structural deviation, exhibited the highest protein expression, with a mean fluorescence intensity (MFI) significantly greater than that of the other constructs (Fig. 4b). Pairwise base-pairing visualization further revealed that CR3 predominantly featured intra-region base pairing within the IRES (blue lines), with minimal inter-region interactions (red lines) (Fig. 4c). In contrast, constructs with higher IRES structural deviation displayed increased inter-region base pairing, which disrupted IRES folding and correlated strongly with diminished protein expression. This observation underscores the essential role of IRES structural integrity in facilitating efficient circRNA translation, likely by ensuring effective recruitment of translation machinery for the translation process.

**Figure 4.**
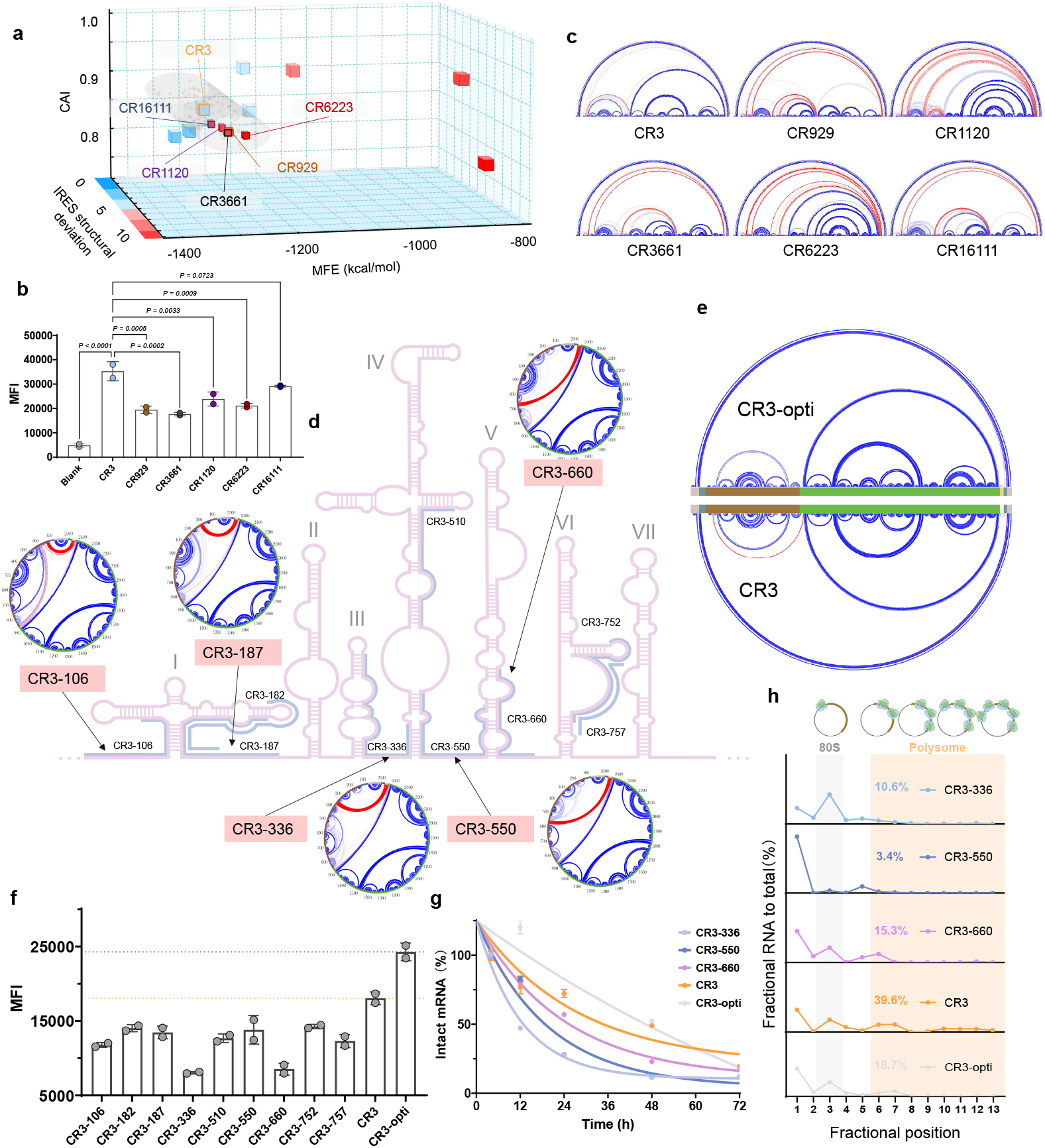
Investigation of the impact of IRES structural deviation on circRNA performance. **a**, Three-dimensional scatter plot illustrating CDS candidates with similar MFE and CAI but varying IRES structural deviation compared to CR3. **b**, Flow cytometry analysis results of protein expression in HSkMC cells 24 hours post-transfection with circRNAs. MFI data are presented as mean ± s.d. from two independent experiments. Statistical analysis was performed using one-way ANOVA with multiple comparisons against the CR3 group. **c**, Pairwise base-pairing visualization of circRNAs CR3, CR929, CR1120, CR3661, CR6223, and CR16111, highlighting varying degrees of IRES folding disruption. Intra-region base pairing is depicted in blue, and inter-region base pairing in red. **d**, Schematic representation of IRES folding disruption walk-through strategy achieved by incorporating sequences complementary to specific IRES regions. The pink RNA secondary structure illustrates the independently folded IRES. Blue lines denote targeted disruption sites within the IRES sequence. Inset: Pairwise base-pairing visualization of CR3-106, CR3-187, CR3-336, CR3-550, and CR3-660, showing progressive disruption of IRES folding. Intra-region base pairing is shown in blue, inter-region in red, with IRES region denoted in brown and CDS in green. **e**, Comparative pairwise base-pairing visualization of CR3 and CR3-opti (displaying base pairs with probabilities ≥ 0.5), where CR3-opti preserves stable IRES folding without directional disruption. Intra-region base pairing is shown in blue, inter-region in red, with IRES region denoted in brown and CDS in green. **f**, Flow cytometry analysis of protein expression in HSkMC cells 24 hours post-transfection with circRNAs designed with specific IRES disruption. MFI values reflect corresponding protein expression levels. Data are presented as mean ± s.d. from two independent experiments. **g**, In-cell degradation kinetics of circRNAs (CR3, CR929, CR1120, CR3661, CR6223, CR16111) post-transfection into cells. circRNA stability was assessed over the course of 72 h, with data presented as mean ± s.d. from four replicates and fitted to a one-phase exponential decay curve. **h**, Polysome profiling analysis of circRNA (CR3-336, CR3-550, CR3-660, CR3, and CR3-opti) across sucrose gradient fractions following ultracentrifugation. The 80S monosome fractions are shaded in gray, and polysome fractions are shaded in orange. Data represent circRNA abundance normalized to total RNA, expressed as mean ± s.d. from three independent biological replicates (*n* = 3).

To further validate the importance of intact IRES folding, we employed a walk-through disruption strategy informed by the secondary structure of the Coxsackievirus B3 (CVB3) IRES (Fig. 4d). We selected distinct 30-nucleotide (nt) functional domains within the IRES and introduced complementary sequences at the junction between CDS and spacer in the circRNA, promoting strong inter-region base pairing intended to disrupt specific IRES regions (Fig. 4d, inset). This approach yielded constructs CR3-106, CR3-182, CR3-187, CR3-336, CR3-510, CR3-550, CR3-660, CR3-752 and CR3-757, which were subsequently compared against both the CR3 control and an optimized variant, CR3-opti, generated by incorporating 30-nt non-complementary sequences at the same junction that mitigate potential disruptions to the IRES, thereby improving its structural integrity. Pairwise base-pairing analysis confirmed that CR3-opti exhibited even lower IRES structural deviation than CR3 (Fig. 4e). Following transfection into HSkMC cells, flow cytometry analysis revealed that CR3-opti achieved the highest protein expression, followed by CR3, while the disrupted constructs displayed reduced protein expression to a varying extent (Fig. 4f). Notably, disruptions in CR3-336 and CR3-660 led to the most significant reductions in expression, whereas CR3-550 showed a milder decrease. These differential effects indicate a hierarchy of functional importance within the IRES, with the regions targeted in CR3-336 and CR3-660 likely playing critical roles in recruiting translation initiation factors or ribosomal subunits, whereas the region disrupted in CR3-550 appears less essential. This targeted disruption approach not only reinforces the importance of IRES structural integrity but also highlights the nuanced contributions of specific domains to circRNA translation efficiency.

We then investigated whether keeping IRES structural integrity extends its intracellular half-life of circRNA. In-cell degradation curve suggested that CR3-opti, with the lowest IRES structural deviation, exhibited the greatest stability (Fig. 4g). CR3 also displayed a slower degradation curve compared with CR3-336, CR-550 and CR-660. Notably, CR3-336, which had lowest protein expression, shown the shortest half-life of approximately 8 hours. This stability trend closely parallels the protein expression data, suggesting that intact IRES folding enhances circRNA longevity, possibly by promoting an actively translating process that reduces the biological decay of circRNA.

To assess the impact from deviated IRES structure on translation process, we performed polysome profiling of selected circRNA followed by reverse transcription quantitative PCR (RT-qPCR) to assess circRNA distribution across different fractions (Fig. 4h). CR3 and CR3-opti exhibited enriched polysome fractions of 39.6 % and 18.7 %, respectively, suggesting robust translation activity. In contrast, constructs such as CR3-660 and CR3-336 showed lower polysome fractions of 15.3 % and 10.6 %, respectively, indicative of reduced translation efficiency. Meanwhile, CR3-550 displayed the lowest polysome fraction at 3.4 %, potentially reflecting a distinct translation profile consistent with its milder expression defect. These findings, combining with protein expression and stability data, corroborated that an integral IRES structure facilitates efficient ribosome engagement and active translation, while disruptions in critical regions hinder this process and thus impact circRNA performance.

Taken together, the correlation observed between IRES integrity, protein output, and RNA stability suggests that a well-folded IRES not only drives efficient translation, but also confers a protective structural advantage that enhances circRNA longevity. These insights have profound implications for the development of circRNA-based therapeutics, where maximizing expression and durability is crucial. By integrating MFE, CAI, and IRES structural deviation into a joint optimization framework, our study provides a blueprint for engineering high-performance circRNAs tailored for biotechnological applications.

## Discussion

In this study, we present circDesign, a novel algorithm for circular RNA (circRNA) sequence generation that aims to enhance circularization efficiency, stability, and translatability. By applying circDesign to the rabies virus glycoprotein (RABV-G) antigen, we demonstrated the vital role of preserving IRES structural integrity in achieving superior circRNA performance. Specifically, when the IRES remains well-folded, circRNAs exhibit greater stability, higher translation efficiency, and, ultimately, stronger *in vivo* immune responses.

A primary limitation of the current work is that our optimization focuses on a single coding sequence (CDS) without further refinement of other critical regions, such as the IRES itself. At present, the *de novo* design of an efficient IRES motif is challenging because virally or cellularly derived IRES elements are generally sensitive to deletions, mutations, or insertions. Their multiple domains must remain properly folded to recruit the translation pre-initiation complex and assemble a functional ribosome. Consequently, we have not yet incorporated direct design of the IRES region in circDesign. Once the structure–function relationships of IRES motifs are characterized at the molecular level, a bottom-up approach to IRES optimization may be integrated into circDesign, potentially enabling fully comprehensive circRNA design.

In addition to IRES-driven translation from circRNA, N^6^-methyladenosine modification upstream of the CDS has been reported to enhance both translation and stability ^31^. Although incorporating chemically modified nucleotides could further broaden the potential of circRNA, we have not yet included these modifications in our current approach. Once explored and validated experimentally, such modifications can be seamlessly integrated to optimize the design process.

Taken together, our findings underscore the promise of circDesign as a robust approach for next-generation vaccine and therapeutic development. By systematically integrating folding stability, codon usage, and IRES integrity into a single framework, circDesign provides a blueprint for generating high-performance circRNA candidates with broad applicability.

## Methods

### Overall algorithmic rationale and innovation

**circDesign** aims to integrate three key objectives for circRNA design:

1. **Stability**: Achieve lower minimum free energy (MFE) values to promote stable secondary structures under a circular folding model.
2. **Translational Efficiency**: Attain favorable Codon Adaptation Index (CAI) values for the CDS region, reflecting the host organism’s preferred codon usage.
3. **IRES Structural Deviation**: Reduce disruptive base-pairing interactions between the IRES and CDS, thereby preserving the functional structure required for cap-independent translation.

Because MFE and CAI impose conflicting constraints, these parameters are balanced through a trade-off strategy, with IRES structural deviation serving as an additional constraint to achieve an overall optimal circRNA design. In practice, we formulate these goals as a single objective function (Eq. 1).

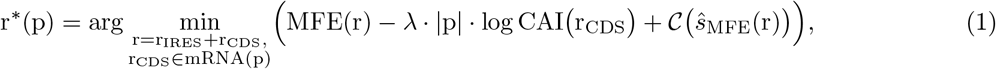

where:

- p: The target protein.
- r = r_IRES_ + r_CDS_: The circRNA sequence consisting of IRES and CDS regions.
- MFE(r): The minimum free energy under an RNA folding model, indicating overall structural stability.
- CAI(r_CDS_): The Codon Adaptation Index, measuring how well the codons in r_CDS_ match the host’s preferred codon usage.
- *λ*: A weighting parameter that balances the stability (MFE) and translational efficiency (CAI).
- *ŝ*_MFE(r)_: The predicted secondary structure corresponding to the minimum free energy (MFE) of sequence r.
- *𝒞 (ŝ*_MFE(r)_) : A structural constraint function that penalizes cross-region base-pairing interactions between IRES and CDS.

By minimizing Eq. 1, **circDesign** identifies sequence variants likely to exhibit favorable stability and expression while maintaining critical IRES functional structure.

We compute the MFE as

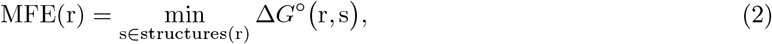

where Δ*G*^*°*^(r, s) is the Gibbs free energy of a secondary structure s for the RNA sequence r. Lower MFE indicates a more stable conformation.

The CAI of an mRNA sequence r that encodes protein p is

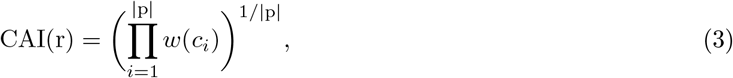

where *w*(*c*_*i*_) represents the relative adaptiveness of codon *c*_*i*_, normalized to the most frequently used synonymous codon in the host organism ^32^. A higher CAI correlates with faster translation rates and potentially increased protein yields.

To help the IRES region retain its functional structure in the circular context, we impose:

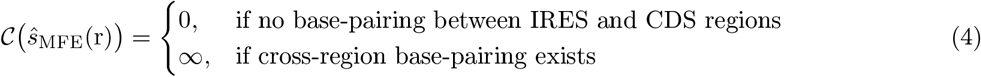

The structural constraint function penalizes sequences with base-pairing interactions between the IRES and CDS, thus reducing the likelihood of IRES structural disruption. Although it does not guarantee the complete absence of cross-region pairing, it significantly lowers the probability of such interference and helps preserve essential IRES motifs for cap-independent translation.

### Beam search and k-best for multi-candidates generation

circDesign employs a beam-search framework, augmented by *k*-best candidate enumeration, to obtain a wide array of candidate sequences with similar key objective parameter values. In particular, when multiple sequences exhibit similarly favorable MFE and CAI values, circDesign further distinguishes them based on cross-region interference, thus favoring those with higher IRES structural integrity.

### *In Silico* screening for further candidate selection

Different circularization strategies may introduce additional elements (such as exons or spacers), or extra sequence fragments may be incorporated for specific purposes. To address this, we implement a post-processing step that calculates the IRES structural deviation, and ensure the structure of IRES retains its integrity within the complete circRNA context. This deviation is defined as the L2 norm between the base-pairing probabilities of the IRES region in a candidate circRNA sequence and those in a reference IRES structure:

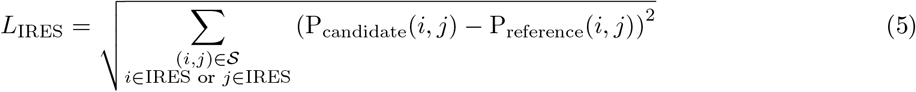

where P(*i, j*) represents the base-pairing probability between nucleotide positions *i* and *j* in an RNA sequence. The candidate probability, P_candidate_(*i, j*), is derived from secondary structure predictions of the entire circRNA sequence, including any additional incorporated elements. In contrast, the reference probability, P_reference_(*i, j*) is obtained from predictions constrained to IRES-internal folding, thereby considering only the self-interactions within the IRES region.

### Additional region for assessing how IRES integrity affects protein expression

To systematically investigate how changes in IRES structural deviation affect protein expression, we introduce an additional 30-nucleotide region downstream of the CDS. This region modulates the IRES structure by:

- Adding sequences complementary to various IRES domains, thereby increasing structural interference.
- Incorporating non-complementary sequences that mitigate potential disruptions to the IRES, thus improving its structural integrity.

### Pseudocodes for circDesign

#### Algorithm 1

CircDesign Algorithm (Nussinov-Based Base-Pairing Rules for Simplification)

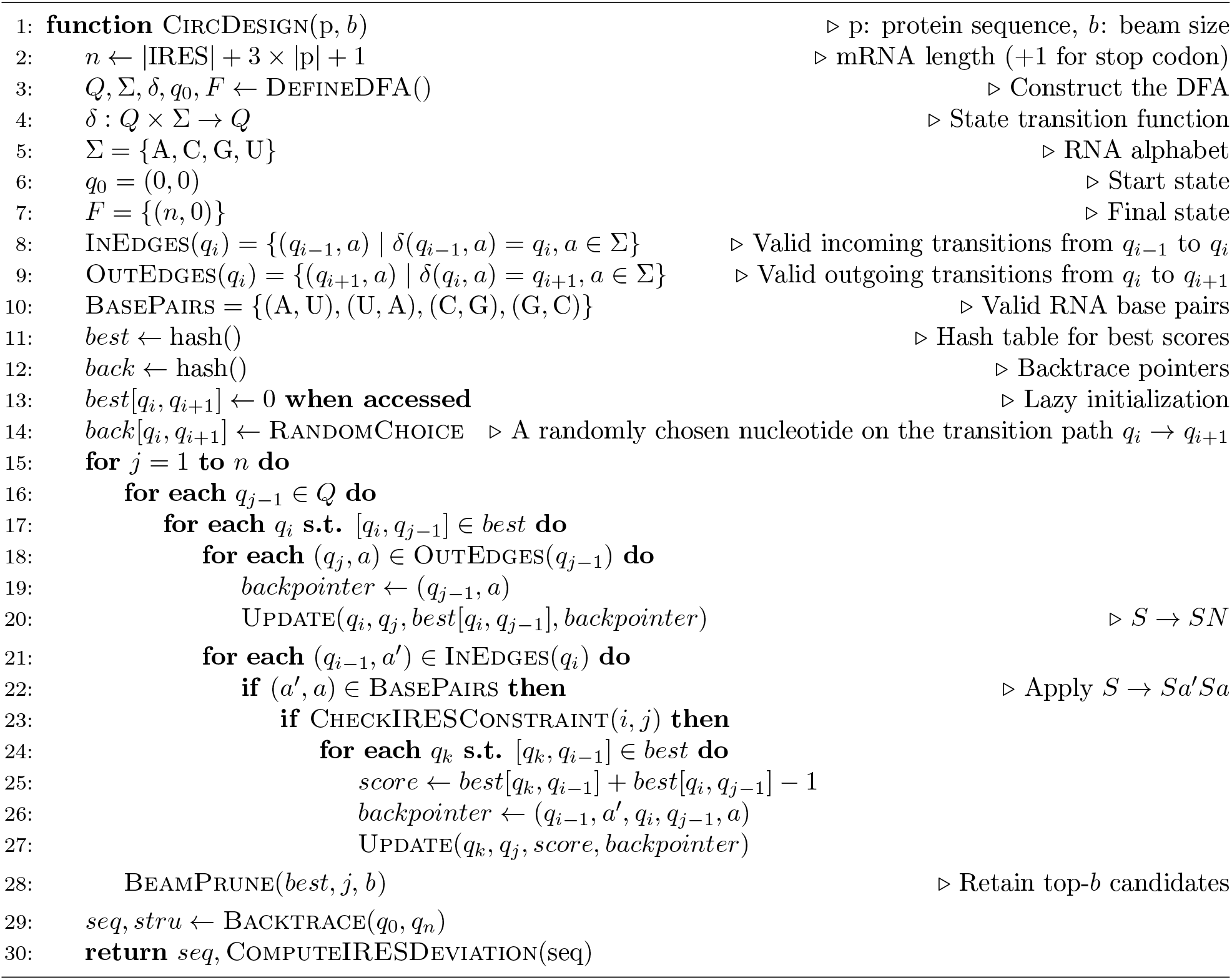

#### Algorithm 2

CheckIRESConstraint Function

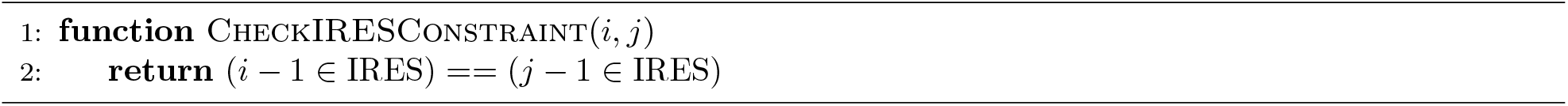

#### Algorithm 3

UpdateIfBetter Function

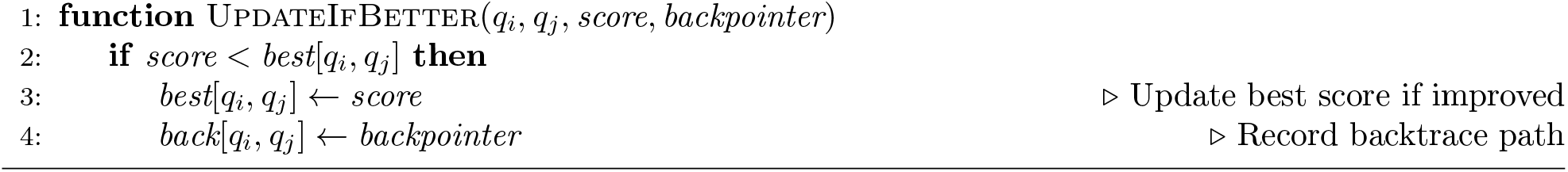

#### Algorithm 4

BackTrace Function

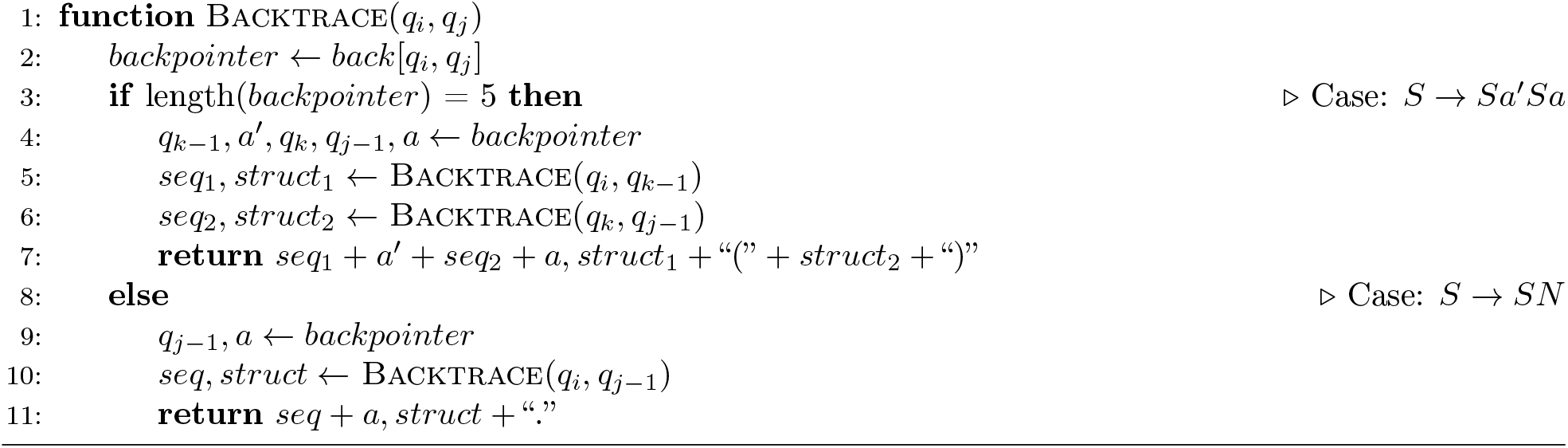

#### Algorithm 5

Compute IRES Structural Deviation

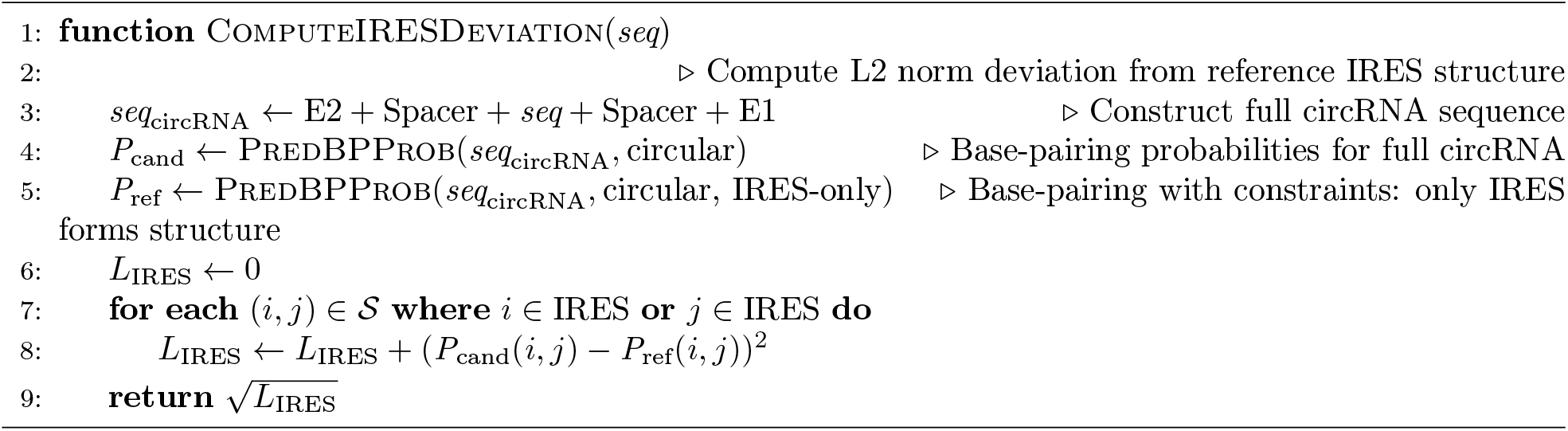

### circRNA secondary structure prediction

The circRNA secondary structure prediction and constraint folding were modeled by ViennaRNA^33,34^. And the structure was visualized by RiboGraphViz and in-house scripts with the cirRNA secondary structure from RNAfold.

### Vector and plasmid construction

To construct a vector for circRNA synthesis, T7 promoter, homology arm, elements from permuted intron-exon (PIE) construct, spacer, IRES, and coding sequences were synthesized by General Biol and cloned into pUC57-mini plasmid. For linear RNA synthesis, the pUC57 plasmid vector containing T7 promoter, 5^*′*^ UTR, Kozak sequence, coding region, 3^*′*^ UTR and polyA tail was synthesized by General Biol. All the sequences coding Firefly Luciferase (FLuc), and RABV-G are supplied in supplemental materials.

### Plasmid DNA synthesis and linearization

pDNA was transformed into *E. coli* (Tsingke Biotechnology) and cultured in 100 mL of Luria Broth (LB) with 100 mg mL^*−*1^ Ampicillin (Sangon Biotech, CN). Plasmid was purified using a NucleoBond Xtra Midi kit (MN) and the concentration and purity was measured on a NanoDrop One (Thermo Fisher). pDNA was linearized using SapI restriction enzyme for 3 h at 37 ^*°*^C and purified as template. After sequencing verification, the products were stored at *−*20 ^*°*^C and were used as templates for *in vitro* transcription (IVT).

### *In vitro* transcription and circularization of RNA

For the chemically modified linear mRNA synthesis, *in vitro* transcription reaction was conducted at 30 ^*°*^C for 16 hours with 100 % N^1^-methylpseudouridine (Hongene Biotech) substitution. mRNA was co-transcriptionally capped using *m*^7^(3^*′*^OMeG)(5^*′*^)ppp(5^*′*^)(2^*′*^OMeA)pG capping reagent (Hongene Biotech) in a “one-pot” reaction followed by digestion with DNase I (Thermo Fisher). Linear mRNA was purified using Dynabeads MyOne Carboxylic Acid beads (Thermo Fisher).

For the circular RNA synthesis, the linear RNA precursor transcripts were produced using 1 µg of linearized DNA template in a MEGAScript^−^ reaction (Ambion) for 16 h at 30 ^*°*^C, according to the manufacturer’s protocol. Then RNA was treated with DNase I (Thermo Fisher) for 60 min at 30 ^*°*^C, followed by lithium chloride precipitation. To obtain circularized RNA, GTP was added to a final concentration of 2 mM along with a buffer including magnesium (50 mM Tris-HCl, (pH 8.0), 10 mM MgCl_2_, 1 mM DTT; Thermo Fisher). RNA was then heated to 55 ^*°*^C and held for 45 min before purification using Dynabeads MyOne Carboxylic Acid beads (Thermo Fisher).

### Liquid chromatography purification of circRNA

Circular RNA products were further purified *via* liquid chromatography to remove spliced introns and precursor linear mRNA using a 21.2 mm × 300 mm size-exclusion column (SEC) with a particle size of 5 µm and pore size of 1000 Å (Sepax Technologies). RNA was run on an AKTA avant 150 purifier system in RNase-free 150 mM phosphate buffer (pH=6.0). Purified circRNA products were concentrated by lithium chloride precipitation and analyzed using Agilent 2100 bioanalyzer.

### CircRNA sequence identification

To validate the circularization of RNA, the SEC purified RNA was reverse transcribed into cDNA using a HiScript III All-in-one RT SuperMix Perfect kit (Vazyme), followed by PCR with primers that can amplify transcripts across the splicing junction site. The PCR products were sequenced using Sanger sequencing to validate the back-splicing junction of the circular RNA.

### Circularization efficiency comparison

To compare the circularization efficiency after circularizing reaction before SEC purification, two sets of primers were used in qPCR method. One set of primers was designed at 3^*′*^ intron, representing the part of RNAs that without successful circularization ; the other set of primers was designed at the junction site, representing the circRNAs formed by successful circularization. Therefore, the circularization efficiency of different RNAs was obtained by comparison, and the sequence of primers was as follows:

- circ-Rabies-F (5^*′*^-3^*′*^): ACAAACGGCTATTATGCGTTACC
- circ-Rabies-R (5^*′*^-3^*′*^): GACTTGAACCCACACGACCG
- Linear-Rabies-F (5^*′*^-3^*′*^):AACGTCAAGACGAGGGTAAAGAGAG
- Linear-Rabies-R (5^*′*^-3^*′*^):GACTTGAACCCACACGACC To calculate the Circularization efficiency of each sequence:

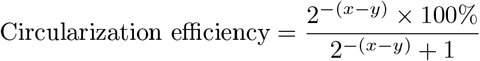

where *x* is the threshold cycle values(Ct) of circular RNA and *y* is the Ct of linear RNA.

### Cell culture and circRNA transfection

Human embryonic kidney 293 (HEK293, ATCC)/Human Skeletal Muscle Cells (HSkMC, ATCC) and Baby hamster kidney 21 (BHK-21, ATCC) were cultured in complete Dulbecco’s Modified Eagle’s Medium (DMEM, Gibco) containing 10 % fetal bovine serum (FBS, Gibco) and 1 % penicillin-streptomycin (Thermo Fisher). Cells were seeded in a 6-well plate at a density of 5 × 10^6^ cells per well 24 h prior to transfection. Lipofectamine Messenger MAX (Thermo Fisher) was used according to the manufacturer’s instructions for the transfection of circRNA.

### Agarose gel electrophoresis of circRNA

To study the electrophoretic mobility profile of circRNA, RNA samples suspended in Ambion^®^ RNA storage buffer (Thermo Fisher) were denatured at 75 ^*°*^C for 5 min and snap-cooled on ice before loaded onto 1 % agrose gel (130 V for 1 h at room temperature). DL5000 DNA marker (Vazyme) was used in the gel. The gel image was taken by Gel Doc XR^+^ Gel Documentation System (Bio-Rad).

### In-solution stability assay of circRNA

To assess the In-solution stability of mRNA, samples were incubated in PBS buffer containing 10 mM Mg^2+^. Sampling was conducted at different time points (0, 1, 2, 4, 8, 12, 16, 24, 32, 48 and 60 hours). RNA integrity was analyzed by Qsep100^−^ Capillary Electrophoresis System (BiOptic Inc). The integrity was represented as the proportion of full-length mRNA calculated on electropherogram. The data were normalized to time point 0 h.

To extrapolate the half-life of each sequence, one-phase decay equation:

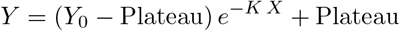

was used to fit the curve. The *Y*_0_ and Plateau were set as 100 and 0, respectively.

The half-life was computed as:

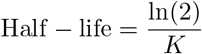

where *K* refers to the decay rate constant.

### Thermodynamic stability assay of circular RNA

Equal volume of circRNA (1 mg mL^*−*1^) and diluted Quant-iT^™^ RiboGreen^®^ (Thermo Fisher) reagent was mixed in 1 × TE buffer. A total of 20 µL mixture was loaded into 384-well plate and centrifuged at 1000 rpm for 1 min before running the Melting Curve program on QuantStudio 5 Real-Time PCR System (Applied Biosystems). circRNA underwent a denaturing process at 95 ^*°*^C for 5 min with a ramp rate of 0.05 ^*°*^C per second starting from 25 ^*°*^C. Then the temperature stepped down to 25 ^*°*^C with a rate of 1.6 ^*°*^C per second for annealing. The fluorescent intensity data at different temperatures were normalized to the initial value and plotted in GraphPad Prism. A sigmoidal curve was used to fit the data to extrapolate the melting temperature (*T*_m_) of each circRNA.

### In-cell stability assay of circular RNA

HSkMC cells were seeded in a 6-well plate at a density of 5 10^6^ cells per well 24 h prior to transfection. Lipofectamine Messenger MAX (Thermo Fisher Scientific) was used according to the manufacturer’s instructions for the transfection of circRNA. Cells were collected and RNA was harvested and purified at 3, 12, 24, 48 and 72 h after transfection using a RNeasy Mini Plus kit (QIAGEN). Synthesis of first-strand cDNA from total RNA was performed with HiScript^®^ III All-in-one RT SuperMix Perfect for qPCR (Vazyme). Gene specific TaqMan primers (TsingKe) amplifying the circRNA junction site were synthesized as below:

- circ-tzF (5^*′*^-3^*′*^): ACAAACGGCTATTATGCGTTACC;
- circ-tzR (5^*′*^-3^*′*^): GACTTGAACCCACACGACCG;
- circ-Probe (5^*′*^-3^*′*^): FAM-ACGGACTTAAAATCCGTTGAC-MGB;

Target amplification sequence was inserted into plasmid as an internal standard to obtain the absolute circRNA copies from RNA extracts. The qPCR reaction was carried out using Probe qPCR Mix kit (Takara). For each sample, threshold cycle values (Ct) were processed according to the comparative Ct method. Gene expression levels were presented relative to Standard curve.

### *In vitro* innate immunogenicity assay

24 hours post transfection of circRNA into C2C12 cells (ATCC), cell culture supernatant was collected. Interferon Beta level in supernatant was assayed by VeriKine−Mouse IFN Beta ELISA Kit (PBL assay science) according to manufacurer’s instruction. Optical density (OD) was measured at 450 nm using Bio-Tek Synergy I plate reader. The concentration of IFN-*β* was deduced from the standard curve.

### Flow cytometry

24 hours post transfection of circRNA, HSkMC cells were harvested and resuspended in 1 mL of PBS buffer at a concentration of 1 × 10^7^ cells*/*mL. 100 µL of the resuspended cells was added to a FACS tube and stained with 100 µL of Live/Dead Fixable Aqua Dead Cell Stain (Thermo Fisher Scientific) at a 1:100 dilution and human FcR blocking reagent (Miltenyi Biotec) at a 1:20 dilution on ice for 30 min. Cells were then washed with 2.5 mL of FACS buffer and centrifuged at 1500 rpm for 5 min. After centrifugation, cells were stained with 1 µg (1:100 dilution) of a anti-Rabies Virus CVS-11 antibody (Merck) at 4 ^*°*^C for 30 min in dark before washing with 2.5 mL of FACS buffer and centrifuging at 1500 rpm for 5 min. Cells were then stained with 100 µL of Goat PE-anti-mouse IgG (1:100 dilution) (Abcam) at 4 ^*°*^C for 30 min in dark. After incubation, cells were washed with 2.5 mL of FACS buffer, centrifuged at 1,500 rpm for 5 min and resuspended with 250 µL of PBS. Samples were analyzed on BD Canto II (BD Biosciences). Data were processed using FlowJo V10.1 (Tree Star).

### RNA sequencing and ribosome profiling of circular RNA

At 6 h post-transfection of CR-Novo and CR3 into HEK293 cells, cycloheximide (Sigma) was added to cell culture medium to a final concentration of 100 µg mL^*−*1^ to arrest protein synthesis and preserve ribosome-mRNA interaction. After a 2-min incubation at 4 ^*°*^C and lysis, the cell lysate was centrifuged at 3000 × g and the supernatant containing free polysomes was decanted. To prepare ribosome footprints (RFs), RNase I (NEB) and DNase I (NEB) were added to lysate and was then incubated at room temperature on a Nutator mixer. Nuclease digestion was stopped by adding SUPERaseIn RNase inhibitor (Ambion) followed by purification using size exclusion columns (GE Healthcare). Next, 10 % (wt/vol) SDS was added to the elution, and RFs with a size greater than 17nt was isolated according to the RNA Clean and Concentrator-25 kit (Zymo Research). After obtaining ribosome footprints, Ribo-seq libraries were constructed using NEBNext^®^ Multiple Small RNA Library Prep Set for Illumina^®^ (catalog no. E7300S, E7300L). Briefly, adapters were added to both ends of RFs, followed by reverse transcription and PCR amplification. The 140– 160bp size PCR products were enriched to generate a cDNA library and sequenced using Illumina HiSeq^−^ X10 by Gene Denovo Biotechnology Co. (Guangzhou, China).

### Ribosome footprint data processing and alignment

Low quality reads were filtered by fastp. Raw reads containing over 50 % of low-quality bases or over 10 % of N bases were removed. Adapter sequences were trimmed. Reads with length between 20–40 bp were retained for subsequent analysis. Short reads alignment tool Bowtie2 was used for mapping reads to ribosome RNA (rRNA) database, GenBank, Rfam database. The reads mapped to rRNAs, transfer RNAs (tRNA), small nuclear RNAs (snRNA), small nucleolar RNAs (snoRNA), and miRNA were removed. Processed RNA-seq reads were aligned to the circular RNA using sequence alignment by STAR with 2-pass setting enabled. The reads were assigned to different functional regions (IRES, CDS, spacer, homology arm and exon) based on the position of the alignment. Translation efficiency (TE) was calculated as the base 2 logarithmic ratio of normalized RPFs over the normalized input for each CDS.

### Pause sites analysis

Because the translocation of ribosomes occurs along the mRNA one codon at a time, the potential of single-codon resolution pauses were investigated. Using PausePred (https://pausepred.ucc.ie/) which infers the locations of ribosome pauses, global single codon pauses were analyzed.

### Quantification of read abundance

Reads counts in the different regions of circular RNA was calculated by software RSEM, and the level was normalized by using FPKM (fragment per kilobase of transcript per million mapped reads) method, and the formula is shown as follows:

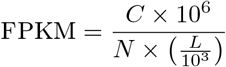

Given FPKM (A) to be the abundance of sequence A, C to be number of fragments mapped to sequence A, N to be total number of fragments that mapped to reference sequences, and L to be number of bases on sequence A. Therefore, the calculated FPKM can be directly used for comparing the difference of abundance among samples. The ribosome footprint depth was plotted in a circular form with each functional region denoted in different color.

### Polysome profiling of circRNA

At 6 h post-transfection of CR-Novo and CR3 into HEK293 cells, cycloheximide (Sigma) was added to cell culture medium to a final concentration of 100 µg mL^*−*1^ to arrest protein synthesis and preserve ribosome-mRNA interaction, and the cells were incubated at 37 ^*°*^C for 15 min. After trypsin digestion (0.25 %), harvest and wash cells with ice-cold PBS supplemented with 100 mg mL^*−*1^ cycloheximide. The frozen tubes were flash frozen in liquid nitrogen and then stored at − 80 ^*°*^C for later use. Samples were sent to NKY-GuangZhou Gene reader Bio-Medical Technologies Co., Ltd. (Guangzhou, China) for isolation. In brief, cells were lysed in lysis buffer (10 mM Tris-HCl pH 7.4, 150 mM NaCl, 5 mM MgCl_2_, 1 % Triton X-100, 0.5 % deoxycholate, 2 mM DTT) supplemented with 4 U mL^*−*1^ RNase inhibitor, 100 µg mL^*−*1^ cycloheximide, and proteinase inhibitor cocktail (Roche, Basel, Switzerland) on ice for 20 min. After centrifugation at 13 000 g for 10 min, the supernatant was then taken and loaded into a 10 % to 45 % sucrose gradient solution and centrifuged at 3.6 × 10^5^ rpm for 3 h (Beckman Optima XE-100). 18 fractions from each sample were collected using a piston gradient fractionator (Biocomp, Fredericton, Canada) equipped with a Bio-Rad Econo UV monitor, followed by total RNA extraction from each fraction for RT-qPCR analysis.

### Preparation of LPP-circRNA

LPP nanoparticles were prepared by a two-step method as previously described. Briefly, a cationic polymer compound (PbAE) was dissolved in PBS buffer solution (pH=5.2) and circRNA was diluted with citrate buffer(pH 4, 5 mM) and RNase-free water. mRNA/PbAE complexes were prepared by a microfluidic mixer (Inano D, Micro&Nano Technology Inc, China) at a volume ratio of 5:1 (mRNA: PbAE). After that, lipids were dissolved in ethanol at molar ratios of 40: 15: 43.5: 1.5 (ionizable lipid: 1,2-dioleoyl-sn-glycero-3-phosphoethanolamine (DOPE): Cholesterol: PEG-lipid). The lipid mixture was combined with mRNA/PbAE complexes at a ratio of 3: 1 (aqueous: ethanol) using a microfluidic mixer. Formulations were concentrated using Amicon ultra centrifugal filters (EMD Millipore) in sodium acetate with 10 % sucrose buffer, passed through a 0.22 mm filter, and stored at 4 ^*°*^C.

### Nanoparticle characterization

The size, polydispersity index (PDI) and zeta potential of LPP formulated circRNA were measured by dynamic light scattering (DLS) (Malvern, Zetasizer Nano ZS). Briefly, LPP-circRNA samples were diluted in sodium acetate with 10 % sucrose buffer at a concentration of 0.1 mg mL^*−*1^. The measurement was performed in triplicate with 8 runs in each measurement.

### *In vitro* storage stability assay of LPP formulated circRNA

LPP formulated circRNA was stored in sodium acetate with 10 % sucrose buffer at 4 ^*°*^C over a time course of 60 days. At Day 0, 30 and 60, LPP-circRNA was sampled for physicochemical characterization including mRNA content, hydrodynamic size and PDI.

### Cell count Kit-8 (CCK-8) assay of LPP formulated circRNA

Cell proliferation capacity was evaluated using CCK-8 assay kit (Sigma Aldrich) according to the manufacturer’s instructions. Briefly, HSkMC cells were seeded into 96-well plates at a density of 5 × 10^5^ cells*/*well. On the other day, 100 ng of LPP formulated circRNA was added into cell culture directly. At 48 h post-transfection, 10 µL CCK-8 solution was added to each well and cells were incubated for a further 2 h at 37 ^*°*^C. Optical density (OD) was measured at 450 nm using Bio-Tek Synergy I plate reader.

### mRNA concentration and encapsulation efficiency of LPP-mRNA

LPP formulated circRNA was diluted with 1 × TE buffer followed by mixing with an equal volume of 2 % Triton X-100 and incubated at RT for 20 min to release the encapsulated mRNA-circRNA core. 100 µL of each sample was then transferred to the wells of a 96-well plate, to which 100 µL of 200 × diluted RiboGreen^−^ RNA Reagent (Thermo Scientific) was added. The plate was shaken and incubated at RT for 10 min. After that, the fluorescence of the plate was read by a Bio-Tek Synergy I plate reader (BioTek). A series of diluted standards were also prepared corresponding to 0.1–2.0 µg*/*mL mRNA. A standard curve of fluorescence as a function of mRNA concentration was plotted by linear regression, from which the mRNA concentrations of the samples were calculated. Encapsulation efficiency was calculated as:

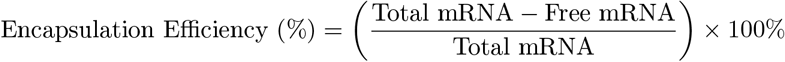

### LPP formulated circRNA protection assay

Content analysis was performed to evaluate the integrity of the RNA after the formulation process and to assess RNase protection of the encapsulated RNA. For the RNase protection assay, 1 mAU of RNase A (Thermo Scientific) per microgram of RNA were added to the LPP/LNP/RNA formulation for 30 min at room temperature. RNase was then inactivated by incubating the sample with 1.68 mAU*/*µg RNA of proteinase K (Thermo Scientific) at 55 ^*°*^C for 10 min. After RNase inactivation, residual RNA was detected from formulations (as described in the **mRNA concentration and encapsulation efficiency of LPP-mRNA** above).

### Fluorescent microscopy imaging of circRNA expression

For the experiments where eGFP-mRNA was transfected, 5 × 10^5^ HSkMC cells were incubated with 2 µg of eGFP-mRNA packaged in LPP or lipofectamine (Thermo Scientific) for 24 h in 6-well plate, and eGFP expression was visualized using an Eclipse DMi8 fluorescent microscope (Leica). Frequency of eGFP-expressing cells was determined using flow cytometry (BD FACSCanto II, Becton Dickinson). The excitation and emission was set at 488 nm and 525 nm, respectively.

### *In vivo* biodistribution of FLuc encoded LPP-circRNAs in mice

Female Balb/c mice aged 6–8 weeks were purchased from Shanghai Model Organisms Center. 5 µg of LPP formulated circRNA or linear RNA encoding Firefly Luciferase (FLuc) in PBS were administrated into mice (*n* = 3 for each group) intramuscularly with 3/10 insulin syringes (BD biosciences). Mice were then anesthetized for whole-body bioluminescence imaging with an IVIS Spectrum (Roper Scientific) at different time points (6 h, 24 h, 48 h, 72 h, 96 h and 192 h).

### Administration of LPP circRNA vaccine in mice

For immunogenicity studies, female Balb/c mice (Shanghai Model Organisms Center) aged 6–8 weeks old (*n* = 5 for each group) were used. 2 µg of LPP formulated circRNA vaccines were diluted in 50 µL × 1 PBS buffer and intramuscularly administrated into the mice’s same hind leg for both prime and boost shots at an interval of 2 weeks. Mice in the control groups received PBS. The blood samples were collected on Day 10, 14, 21 and 28 post prime dosing. Mice spleens were also harvested at the endpoint (D28) for ELISpot and flow cytometry.

### pVNT measurement

A pseudovirus neutralization assay was performed to evaluate the neutralizing ability of sera collected from immunized mice. In brief, serially diluted antibodies were first incubated with pseudotyped virus (Beijing Tiantan Biological) for 1 h, and the mixture was then incubated with BHK-21 cells. After a 24-h incubation in an incubator at 37 ^*°*^C, cells were collected and lysed with luciferase substrate (PerkinElmer), then underwent luminescence intensity measurement by BioTek microplate reader. The IC_50_ value was calculated using 4 parameter logistic non-linear regression.

### *In vivo* mouse cytokine analysis

A serial dosage of LPP formulated CR3, CR-Novo and LR-Novo (0.02 µg, 0.2 µg, 2 µg) were intramuscularly administered into female Balb/c mice (*n* = 5, Shanghai Model Organisms Center) aged 6–8 weeks old. Mice sera were collected retro-orbitally at 4 h and 7 d post administration and subjected to cytokine analysis (Cytometric Bead Array Mouse Inflammation Kit, BD) using flow cytometry following the manufacturer’s instructions. Data were presented as the percentage change relative to baseline mice group administered with PBS.

### IFN-*γ* ELISpots

Assessment of the IFN-*γ* T cell response was performed using the Mouse IFN-*γ* ELISpotPLUS kit (Mabtech) following the manufacturer’s instructions. Briefly, anti-IFN-*γ* pre-coated plates were blocked with DMEM+10 % FBS for at least 30 min, then cells were added at 3× 10^5^ cells per well for negative control (media only) and RABV-G peptide pools (15-mers overlapping by 11; Genscript) (1 µg mL^*−*1^) in 200 µL final volume per well. The positive control wells contained 5 × 10^4^ cells per well in 200 µL final volume per well with 5 µg mL^*−*1^ of ConA. Plates were incubated overnight at 5 % CO_2_, 37 ^*°*^C incubator and developed as per the manufacturer’s protocol. After overnight stimulation, plates were washed and sequentially incubated with biotinylated IFN-*γ* detection antibody (R4-6A2), streptavidin-ALP, and finally BCIP/NBT. Plates were imaged with ImmunoSpot Analyzer and quantified with ImmunoSpot software.

### Biodistribution of LPP-circRNA

The mice were divided into two groups, and 5 µg CR-Novo-LPP or CR3-LPP was injected into the hind legs, respectively. 5 mice in each group were euthanized at 2 h, 6 h, 24 h, 48 h, 72 h and 168 h after administration. Tissue samples including heart, liver, spleen, lung, kidney, small intestine, brain, muscle at the injection site, and lymph node were collected and rapidly frozen in liquid nitrogen, and RNA was extracted using trizol method. The levels of CR-Novo or CR3 in tissue samples were analyzed by RT-qPCR. GraphPad Prism 8 and Origin2021 software were used for data processing.

Gene specific TaqMan primers amplifying the circRNA junction site were synthesized (TsingKe) as below:

- circ-tzF (5^*′*^-3^*′*^): ACAAACGGCTATTATGCGTTACC
- circ-tzR (5^*′*^-3^*′*^): GACTTGAACCCACACGACCG
- circ-Probe (5^*′*^-3^*′*^): FAM-ACGGACTTAAAATCCGTTGAC-MGB

Target amplification sequence was inserted into plasmid as an internal standard to obtain the absolute circRNA copies from RNA extracts. The qPCR reaction was carried out using Probe qPCR Mix kit (Takara). For each sample, threshold cycle values (Ct) were processed according to the comparative Ct method. Gene expression levels were presented relative to Standard curve.

### Statistical Analyses

The significance between groups was determined using a one-way ANOVA with Dunn’s multiple comparison test. Prism version 8.0 (GraphPad) or Origin 10.0 were used for generation of all graphs and performance of statistical analyses. Statistical significance is denoted as ns, not significant, * *p <* 0.05, ** *p <* 0.01, and *** *p <* 0.001.

## Data Availability

The circRNA sequences used in the biological experiments are included at the end of the Supplementary Information file and are available at our figshare repository. Source data are provided with this paper.

## Code Availability

Upon acceptance of this manuscript, circDesign will be made publicly accessible as a web server. Detailed pseudocode for the algorithms is provided in the Methods section.

## Acknowledgements

We thank the assistance from StemiRNA Therapeutics colleagues including Xinxin Su, Xuhui Qiu, Heng Xu, Mingyang Liu, Bin Li, Yun Ge and Kun Xie at StemiRNA Therapeutics in sample preparation and animal experiments. This work was supported by the National Key R&D Program of China (2024YFA0916700, 2024YFA0918600, 2021YFB3800900), National Natural Science Foundation of China (52233007, 32470995, 22477122, 32270627), National Basic Research Plan of China (2023YFA0915201, 2024YFA1308500), Beijing Municipal Science & Technology Commission (Z231100007223003), and Beijing Natural Science Foundation (JQ24007). Congcong Xu is sponsored by Shanghai Pujiang Talent Program (22PJ1423100). All the animal experiments were approved by the Animal Care and Use Committee of Soochow University (Suzhou, China), and all protocols of animal studies conformed to the Guide for the Care and Use of Laboratory Animals (approval NO. SYXK 2021-0065).

## Author contribution

C.X. and L.Z. conceived and directed the project. L.Z. and C.X. introduced the key concept of IRES structural deviation in circRNA design algorithm. C.X., L.Z., D.L., H.S., W.T. and Z.Z. provided important resources and supervised the project. C.X., L.Z. and D.L. supervised all the *in vitro* and *in vivo* experiments. L.Z. designed and developed the circDesign algorithm and L.Z., C.X., C.P. and F.J. designed circRNA sequences. C.X., M.L., R.C., W.W., and J.A. performed the mRNA synthesis and gel electrophoresis experiments. W.W., R.C., J.A., D.Z., Y.C., W.H., C.D., Y.Z., X.W., X.S., Y.Y. and H.S. performed the protein expression and *in vivo* assays, and Q.W. conducted LPP formulation and chemical stability assays. C.D., Y.T. and R.C. performed the ribo-seq experiments. H.Z. and Y.G. performed the animal study including injection and *in vivo* imaging. C.X., L.Z., C.P., R.C., H.L., W.W., Y.T., and Q.W. wrote the manuscript. C.X., L.Z., D.L., Z.Z. reviewed and edited the manuscript.

## Ethics declarations

All mouse studies were performed in strict accordance with the guidelines set by the Chinese Regulations of Laboratory Animals and Laboratory Animal-Requirements of Environment and Housing Facilities. Animal experiments were carried out with the approval from the Institutional Animal Care and Use Committee (IACUC) of Shanghai Model Organisms Center and Soochow University (Approval number SYXK 2021-0065).

