## Extended data for "circDesign Algorithm for Designing Synthetic Circular RNA"

### Extended Data Figures

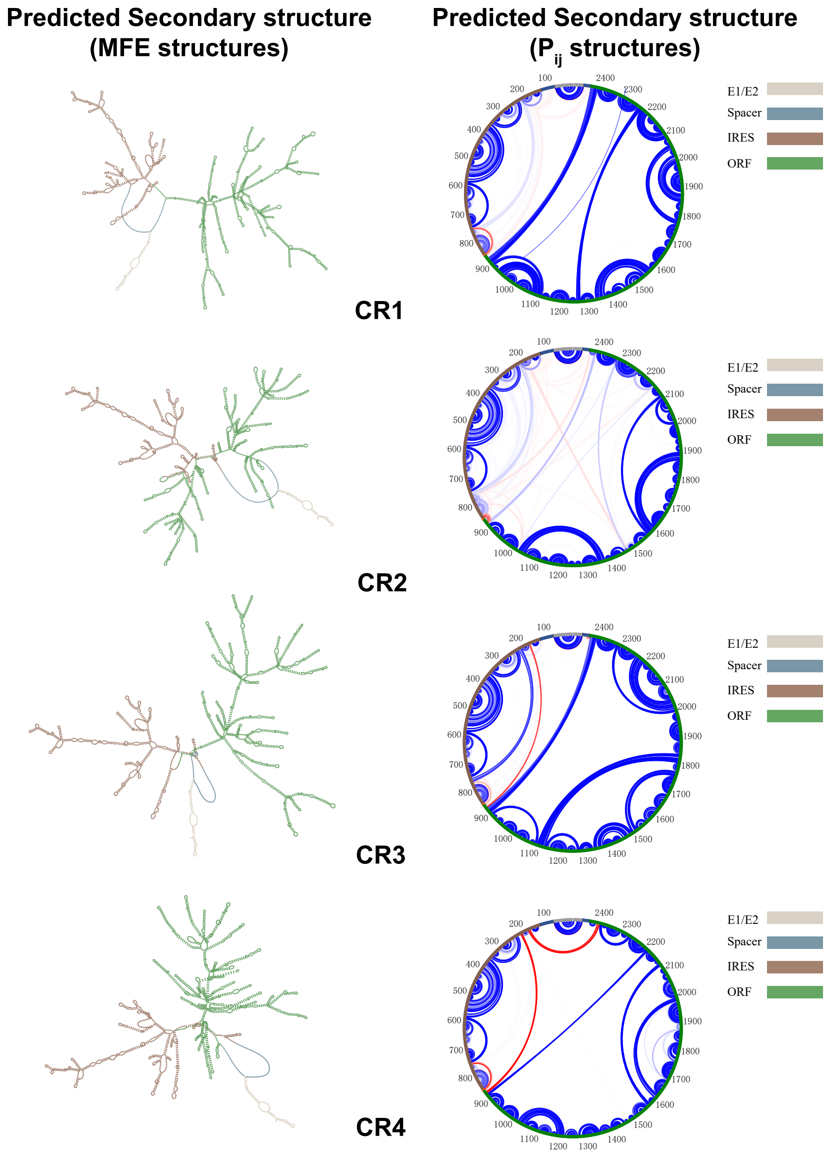


**Extended Data Figure 1. Secondary structure predictions of CR1–CR4.** Left: Predicted minimum free energy (MFE) structures. Right: Base-pairing probability ($P_{ij}$) structures, with intra-region and inter-region base-pairing interactions shown in blue and red. Each region is color-coded as indicated.

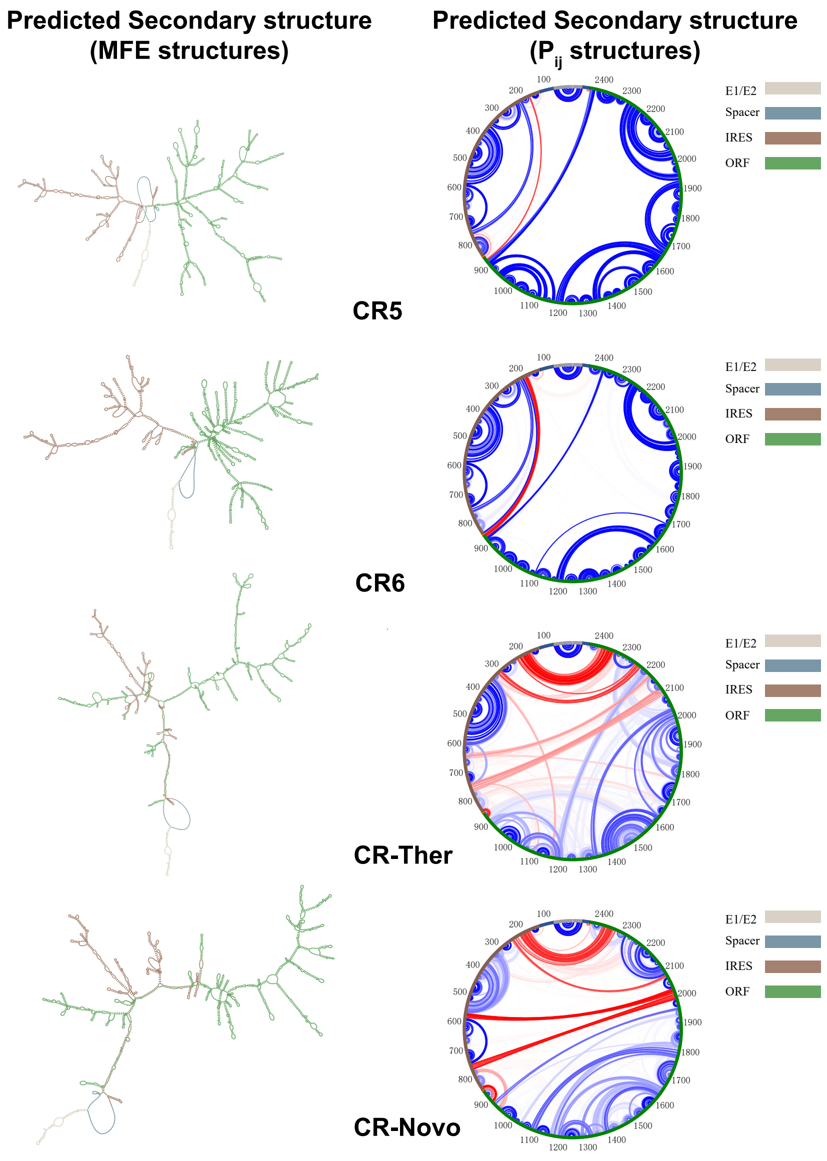


**Extended Data Figure 2. Secondary structure predictions of CR5, CR6, CR-Ther and CR-Novo.** Left: Predicted minimum free energy (MFE) structures. Right: Base-pairing probability ($P_{ij}$) structures, with intra-region and inter-region base-pairing interactions shown in blue and red. Each region is color-coded as indicated.

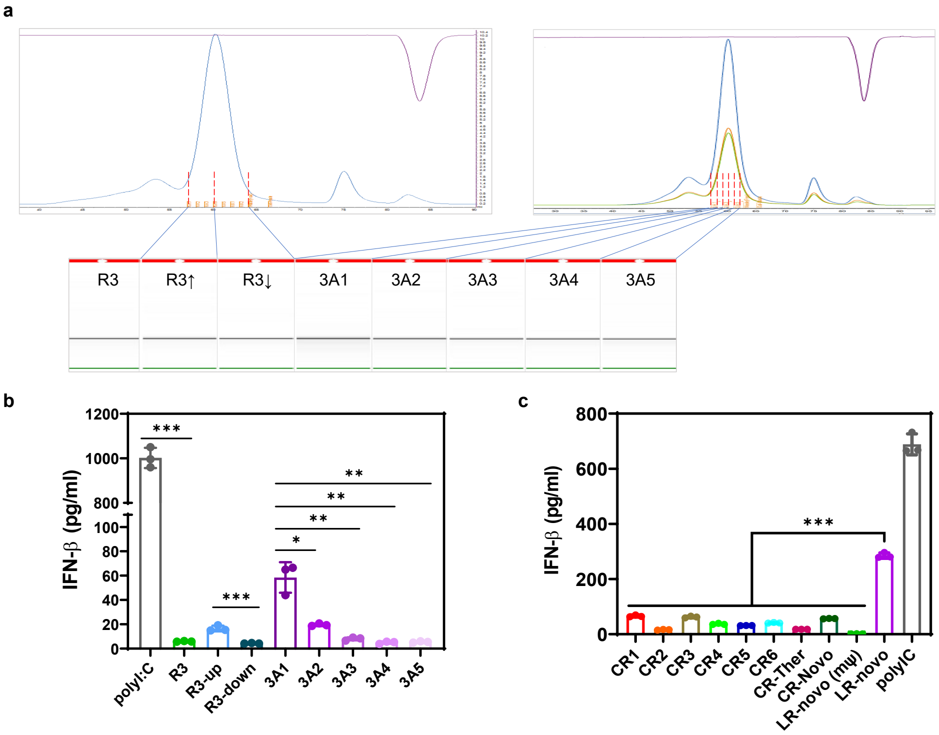


**Extended Data Figure 3. Purification and innate immunogenicity of circRNA. a,** Size-exclusion column (SEC) chromatography purification of circRNA. The major peak was crudely segmented into two fractions as R3↑ and R3↓ (left panel). The major peak was separated into five fractions as 3A1–3A5 (right panel). **b,** *In vitro* interferon $\beta$ expression level from cell culture 24 h post-transfection of circRNAs collected from different SEC fractions. **c,** *In vitro* interferon $\beta$ expression level from cell culture 24 h post-transfection of different SEC purified circRNAs or linear mRNA (LR-Novo, m$\psi$ refers to N1-methylpseudouridine modification).

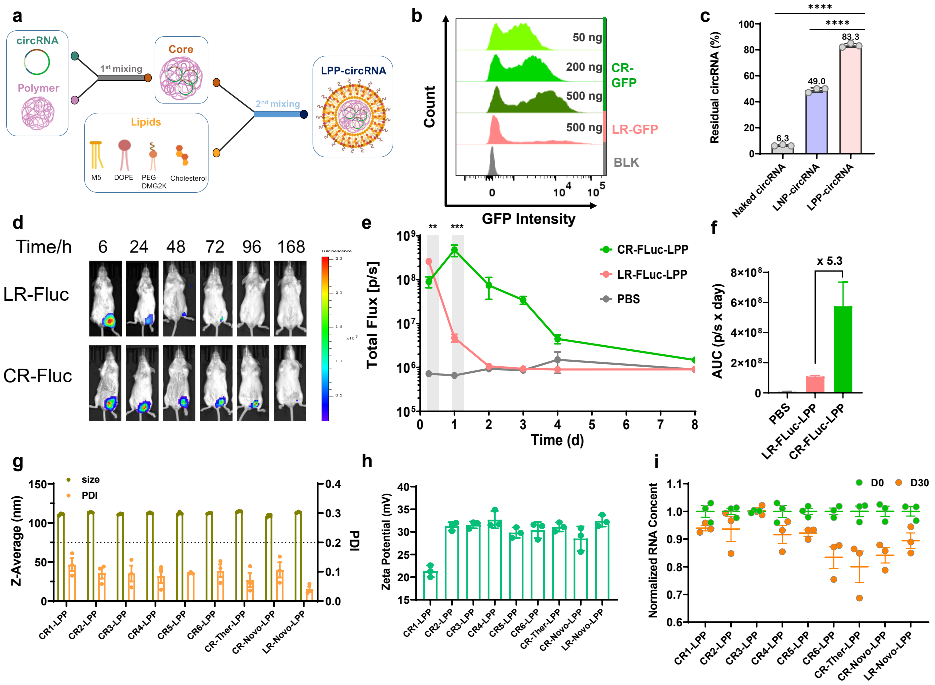


**Extended Data Figure 4. Characterization of Lipopolyplex (LPP)-formulated circRNAs. a,** Schematic representation of the two-step manufacturing process for LPP-formulated circRNA. The lipid mixture consists of an ionizable lipid, a PEGylated lipid, cholesterol, and a helper lipid, combined to encapsulate the circRNA core into a stable nanoparticle structure. **b,** Hydrodynamic size (in nanometers, nm) and polydispersity index (PDI) of LPP-formulated linear RNA (LR-FLuc) and circular RNA (CR-FLuc), determined by dynamic light scattering (DLS). Data are represent as mean ± s.d. from three independent measurements ($n = 3$). **c,** Zeta potential (in millivolts, mV) of LPP-formulated circRNA and linear RNA, reflecting the surface charge properties of the nanoparticles. Measurements were conducted in triplicate, with data presented as mean ± s.d. ($n = 3$). **d,** Representative whole-body *in vivo* imaging system (IVIS) images of Balb/c mice administered LPP-formulated CR-FLuc or LR-FLuc. Bioluminescence was recorded at 6, 24, 48, 72, 96, and 168 hours post-injection to monitor luciferase expression over time. **e,** Total luminescence flux (in photons per second) plotted as a function of time in Balb/c mice ($n = 3$ per group). Data are shown for CR-FLuc-LPP (green), LR-FLuc-LPP (pink), and a phosphate-buffered saline (PBS) control (gray), illustrating the temporal dynamics of luciferase activity. **f,** Area under the curve (AUC) of luminescence from 6 to 168 hours, quantifying the cumulative luciferase activity for CR-FLuc-LPP, LR-FLuc-LPP, and PBS groups. Each data point represents the mean ± s.e.m. from three mice. **g,** Hydrodynamic size (in nm) and PDI of LPP-formulated circRNA and linear RNA, verifying the consistency of nanoparticle size and distribution. Data are presented as mean ± s.d. from three independent measurements ($n = 3$). **h,** Zeta potential (in mV) of LPP-formulated circRNA and linear RNA, further characterizing the surface charge of the nanoparticles. Data are presented as mean ± s.d. from three independent measurements ($n = 3$). **i,** Storage stability of LPP-formulated circRNA and linear RNA at 4 °C, evaluated by assessing RNA integrity over time to infer the shelf-life potential of circRNA vaccines. Samples were tested with RNA integrity normalized to day 0 (set as 1). Data are presented as mean ± s.d. from three independent measurements ($n = 3$).

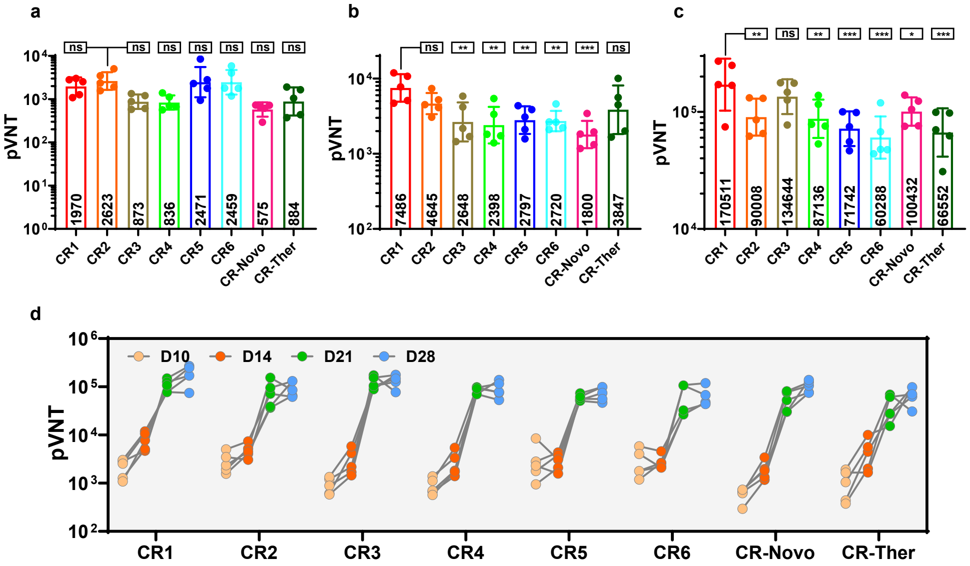


**Extended Data Figure 5.** ***In vivo*** **immune responses induced by circRNA rabies vaccine.** **a–c,** Pseudovirus neutralizing titer (pVNT) of mice sera immunized with two doses of LPP formulated RNA vaccines on D10 (a), D14 (b) and D28 (c). pVNT data were presented as mean ± geometric s.d.. **d,** Summary of pVNT results from D10 to D28 of all vaccinated Balb/c mice ($n = 5$) Data were presented as geometric mean ± geometric s.d..

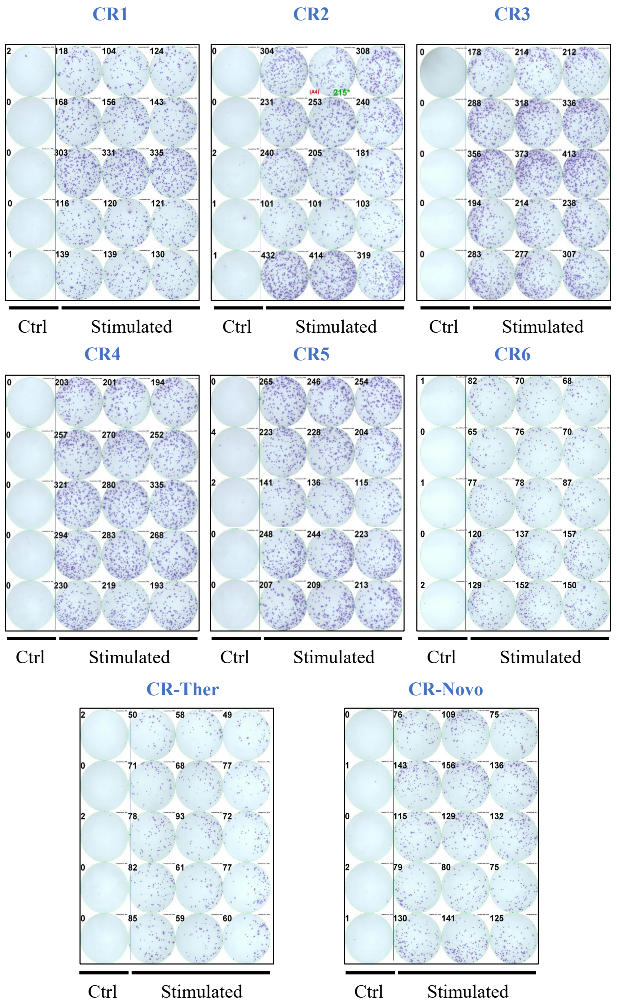


**Extended Data Figure 6. IFN-$\gamma$ ELISpot analysis of mice splenocytes on Day 28.** circDesign-generated sequences show robust cellular immune responses triggered. In each panel, the first column represents control group (unstimulated cells) while the rest three column refers to three technical replicates of peptide pool stimulated sample. Each row represents the sample from one individual mouse.

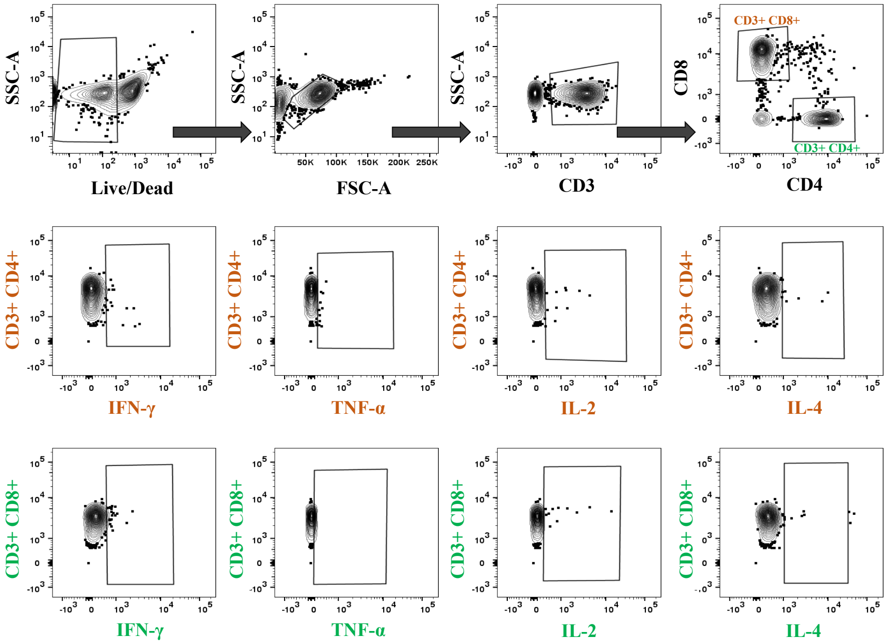


**Extended Data Figure 7. The gating strategy for FACS analysis of cell-mediated immune response in mice vaccinated with circRNA vaccines.** Representative images from an individual mouse splenocyte sample were shown to process the FACS data step-by-step and the same gating strategy was applied to all other experimental groups.

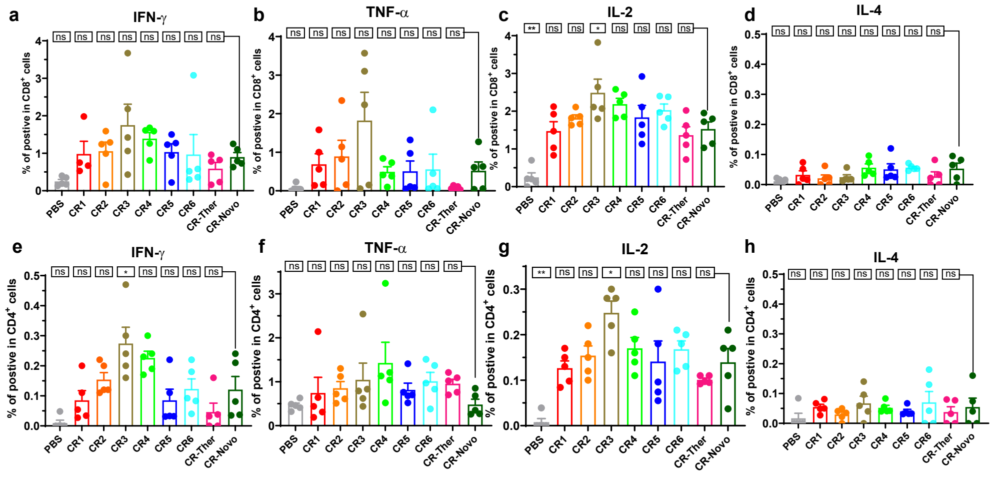


**Extended Data Figure 8.** ***In vivo*** **immune response analysis of LPP-Delivered RNA rabies vaccines. a–d,** Flow cytometry (FACS) analysis of $CD8^+$ T cells secreting IFN-$\gamma$ (a), TNF-$\alpha$ (b), IL-2 (c) and IL-4 (d), following stimulation with a rabies virus glycoprotein (RABV-G) peptide pool. **e–h,** FACS analysis of $CD4^+$ T cells secreting IFN-$\gamma$ (e), TNF-$\alpha$ (f), IL-2 (g) and IL-4 (h), following stimulation with a rabies virus glycoprotein (RABV-G) peptide pool.

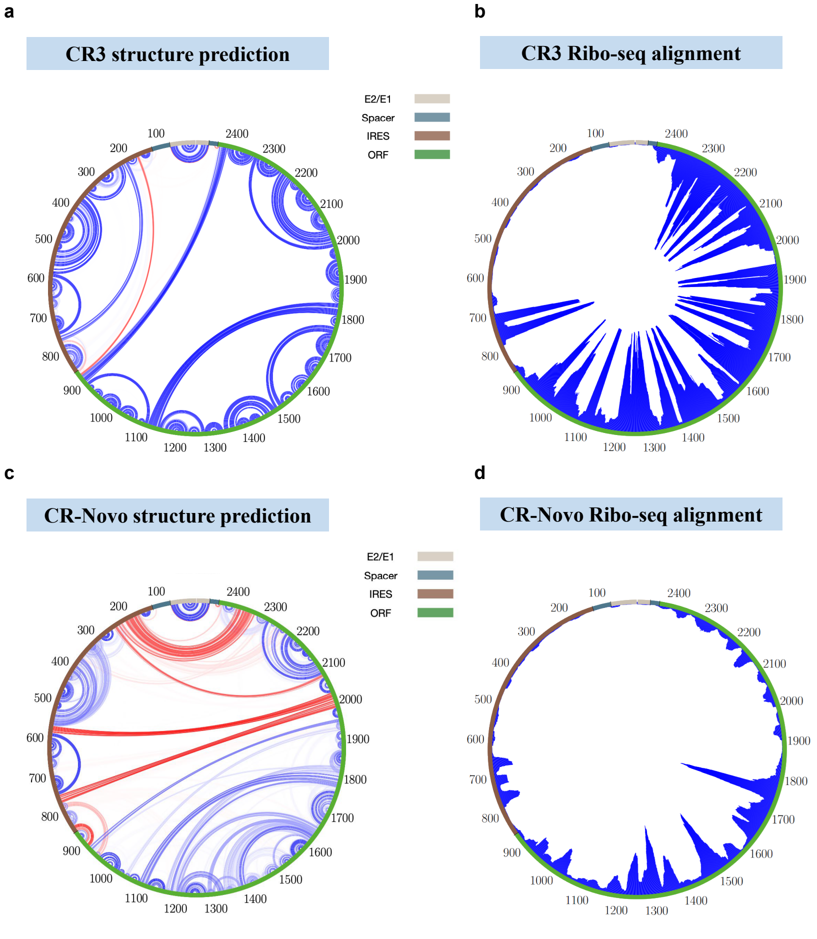


**Extended Data Figure 9. Circularized RNA base pairing probability and ribosome footprints alignment.** Pairwise visualization of base pairing in CR3 (a) and CR-Novo (c). The circular plot showing ribosome footprint depth on CR3 (b) and CR-Novo (d). Each functional region was labeled with different color as indicated.

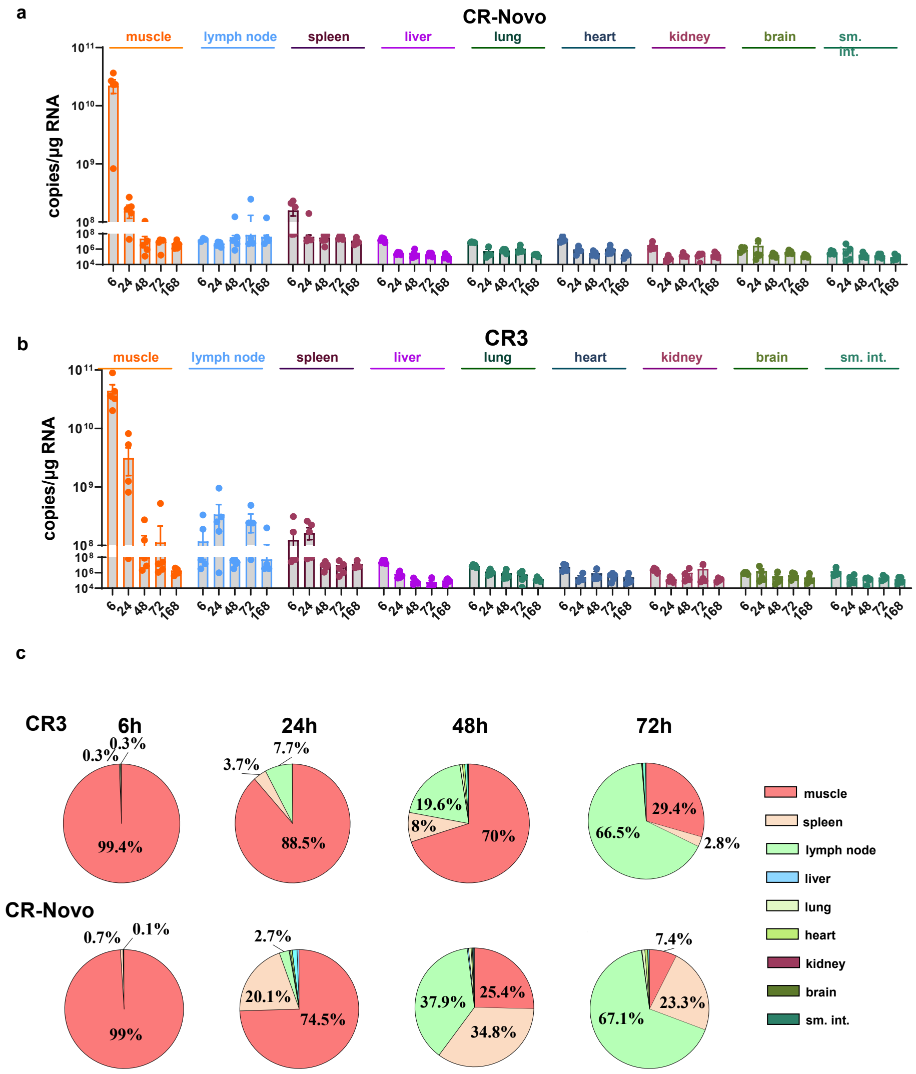


**Extended Data Figure 10.** ***In vivo*** **biodistribution of LPP formulated circRNA. a,b,** RT-qPCR quantification of CR-Novo (a) and CR3 (b) in different tissues at different time points (6 h, 24 h, 48 h, 72 h, 168 h) in Balb/c mice ($n = 5$ per group at each time point). **c,** Pie plot showing the tissue distribution of circular RNA at various time points.
